# Chemotherapy but not the tumor draining lymph nodes determine the immunotherapy response in secondary tumors

**DOI:** 10.1101/664912

**Authors:** Xianda Zhao, Beminet Kassaye, Dechen Wangmo, Emil Lou, Subbaya Subramanian

## Abstract

Immunotherapies are used as adjuvant therapies for cancers. However, knowledge of how traditional cancer treatments affect immunotherapies is limited. Using mouse models, we demonstrate that tumor-draining lymph nodes (TdLNs) are critical for tumor antigen-specific T-cell response. However, removing TdLNs concurrently with established primary tumors did not affect the immune checkpoint blockade (ICB) response on localized secondary tumor due to immunotolerance in TdLNs and distribution of antigen-specific T cells in peripheral lymphatic organs. Notably, treatment response improved with sequential administration of 5-fluorouracil (5-FU) and ICB compared to concurrent administration of ICB with 5-FU. Immune profiling revealed that using 5-FU as induction treatment increased tumor visibility to immune cells, decreased immunosuppressive cells in the tumor microenvironment, and limited chemotherapy-induced T-cell depletion. We show that the effect of traditional cytotoxic treatment, not TdLNs, influences immunotherapy response in localized secondary tumors. We postulate essential considerations for successful immunotherapy strategies in clinical conditions.

**Graphic abstract:** The effects of tumor-draining lymph nodes (TdLNs) resection and a combination of cytotoxic chemotherapy on immune checkpoint blockade therapies are evaluated in this study. TdLNs resection was adverse in eliciting an antitumor immune response in early-stage tumors, but not in late-stage tumors. Further, sequential treatments with cytotoxic chemotherapy and immunotherapy showed better tumor control compared to concurrent combinatorial treatments.

## INTRODUCTION

Immune checkpoint blockade therapies (ICBT) such as anti-CTLA-4 and anti-PD-1/PD-L1, have transformed the therapeutic landscape of cancers, including melanoma and tumors with microsatellite instability (Le et al., 2017; Robert et al., 2011; Topalian et al., 2012). Nonetheless, as with more traditional forms of systemic chemotherapy options, many patients manifest either intrinsic or acquired resistance leading to treatment failure (Gide et al., 2018; Sharma et al., 2017; Zhao and Subramanian, 2017). Multiple mechanisms that influence tumor response to ICBTs have been identified— the mutational load in tumor cells, degree of T-cell exhaustion, tumor microenvironmental functions, and intestinal microbiota (Gide et al., 2018; Sharma et al., 2017; Zhao and Subramanian, 2017). In most cases, ICBT is used for treating patients with heavily pre-treated tumors. The interactions between first-line therapy may influence tumor response to subsequently administered ICBTs due to tumor evolution and heterogeneity. In most patients with solid tumors, common interventions before ICBT include resection of primary tumors with concurrent resection of draining lymph nodes followed by administration of chemotherapies and/ or targeted therapies (Le et al., 2017; Lee et al., 2017; Rizvi et al., 2015). However, minimal information is known about whether these interventions will impact tumor response to ICBT.

Tumor-draining lymph nodes (TdLNs), that are usually resected concurrently with the primary tumors, have shown dual impacts on tumor development and treatment. On the one hand, TdLNs are critical peripheral lymphatic organs where tumor antigens are presented by dendritic cells to naïve T cells to elicit antitumor immunity (Fisher and Fisher, 1971; Shu et al., 2006; Toki et al., 2020). Thus, loss of TdLNs weakens immunosurveillance mechanisms and increases the likelihood of tumor initiation and progression (Fisher and Fisher, 1971; Karlsson et al., 2010; Shu et al., 2006; Toki et al., 2020). On the other hand, TdLNs are affected by immunosuppressive factors released by tumor cells during tumor progression. These immunosuppressive factors can suppress the function of TdLNs, making them immune-privileged sites (Cochran et al., 2001; Ito et al., 2006; Munn and Mellor, 2006; Murthy et al., 2019; Watanabe et al., 2008). Based on these facts, we hypothesize that TdLN resection is an essential factor which influences long-term tumor immunity and response to ICBT. In this study, we used tumor models representing different disease stages to elucidate the impacts of TdLN resection on ICBT efficacy and understand the underlying mechanisms of those effects.

The immunoregulatory effects of chemotherapies have been investigated in multiple cancer models with different chemotherapy drugs. Chemotherapy drugs such as oxaliplatin, paclitaxel, and 5-fluorouracil (5-FU) have shown positive effects in antitumor immunity either by eliciting a tumor-specific T-cell response or by reducing immunosuppressive factors in the tumor microenvironment (Khosravianfar et al., 2018; Pfirschke et al., 2016; Zhang et al., 2008). Bone marrow suppression, which is a common side effect of chemotherapies, causes leukopenia that affects antitumor immunity. Because chemotherapies have dual effects on regulating antitumor immunity, we hypothesize that combining chemotherapy with ICBT has diverse effects on antitumor immune response and consequently, an appropriate combinatory strategy will be critical in determining tumor response. In this study, we used 5-FU, which blocks DNA replication, as a representative chemotherapeutic drug to study the factors that influence the effects of chemotherapy on ICBT.

Mouse models are critical for pre-clinical cancer studies; most published studies have been performed on primary tumor models. To better represent the clinical conditions in which most immunotherapies are administered, we established a mouse tumor model that allows evaluation of the immunotherapeutic response in secondary tumors after primary tumor resection with or without concurrent TdLN removal. We also included anti-PD-1 (antagonist to inhibitory immune checkpoints) and anti-4-1BB (agonist to stimulatory immune checkpoints) to better represent ICBT with different mechanisms (Buchan et al., 2018; Chester et al., 2016).

## RESULTS

### TdLNs are essential for antitumor immune activation and immunotherapy response in early-stage disease

We first needed to identify TdLNs in the subcutaneous tumor model. We injected Evans blue and Alexa Fluor 488 into tumors established in the right flank of the mice to trace lymphatic drainage (Figure S1A). Evans blue staining was detected in the right inguinal and axillary lymph nodes 10 min after injection (Figure S1B). To develop a more sensitive method for detection, we used flow cytometry to trace the Alexa Fluor 488 drainage in lymphatic organs for up to 48 hr. Again, the right inguinal and right axillary lymph nodes showed the highest fluorescence intensity (Figure S1C). Other lymph nodes, such as right brachial and right popliteal lymph nodes, also showed increased fluorescence signal after injection, but the signal intensity was significantly lower than in the right inguinal and right axillary lymph nodes (Figure S1C). Also, increased weight was observed in the spleen, and right inguinal and axillary lymph nodes during tumor development, suggesting an immune response occurred in these lymphatic organs (Figure S2A-J). Collectively, these results indicated that the right inguinal and right axillary lymph nodes are the sentinel TdLNs in our tumor model.

Next, we evaluated the impact of TdLNs on tumor initiation and antitumor immune response stimulation. Resection of TdLNs, but not non-draining lymph nodes (NdLNs), before tumor cell inoculation significantly accelerated tumor development in both CT26 and MC38 tumor models (Figure 1A). We then analyzed the stimulation of antitumor immunity with and without TdLNs. We used the frequency of tumor antigen-specific CD8^+^ T cells (10 days after tumor cells inoculation) as an indicator of antitumor immune response stimulation (Figure S3). More tumor antigen-specific CD8^+^ T cells were detected in the right and left brachial lymph nodes and spleen of tumor-bearing mice with intact TdLNs (Figure 1B). 4-1BB (CD137) provides important co-stimulatory signaling for T cells, and its agonist has shown tumor-eliminating effects in mice (Chester et al., 2016). To test the effects of TdLNs on the immunotherapeutic response, we administered two injections of anti-4-1BB shortly after tumor cell inoculation to simulate patients with minimal disease burden. The prophylactic anti-4-1BB treatments successfully prevented tumor development in mice with intact TdLNs, and anti-tumor immunity memory was established as evidenced by their rejection of secondary tumors. These effects were not seen in mice with resected TdLNs (Figure 1C). These data demonstrated that TdLNs are critical for anti-tumor immunity activation and loss of TdLNs leads to rapid early-stage tumor growth even with the potent T-cell co-stimulatory agonist.

**Figure 1.**
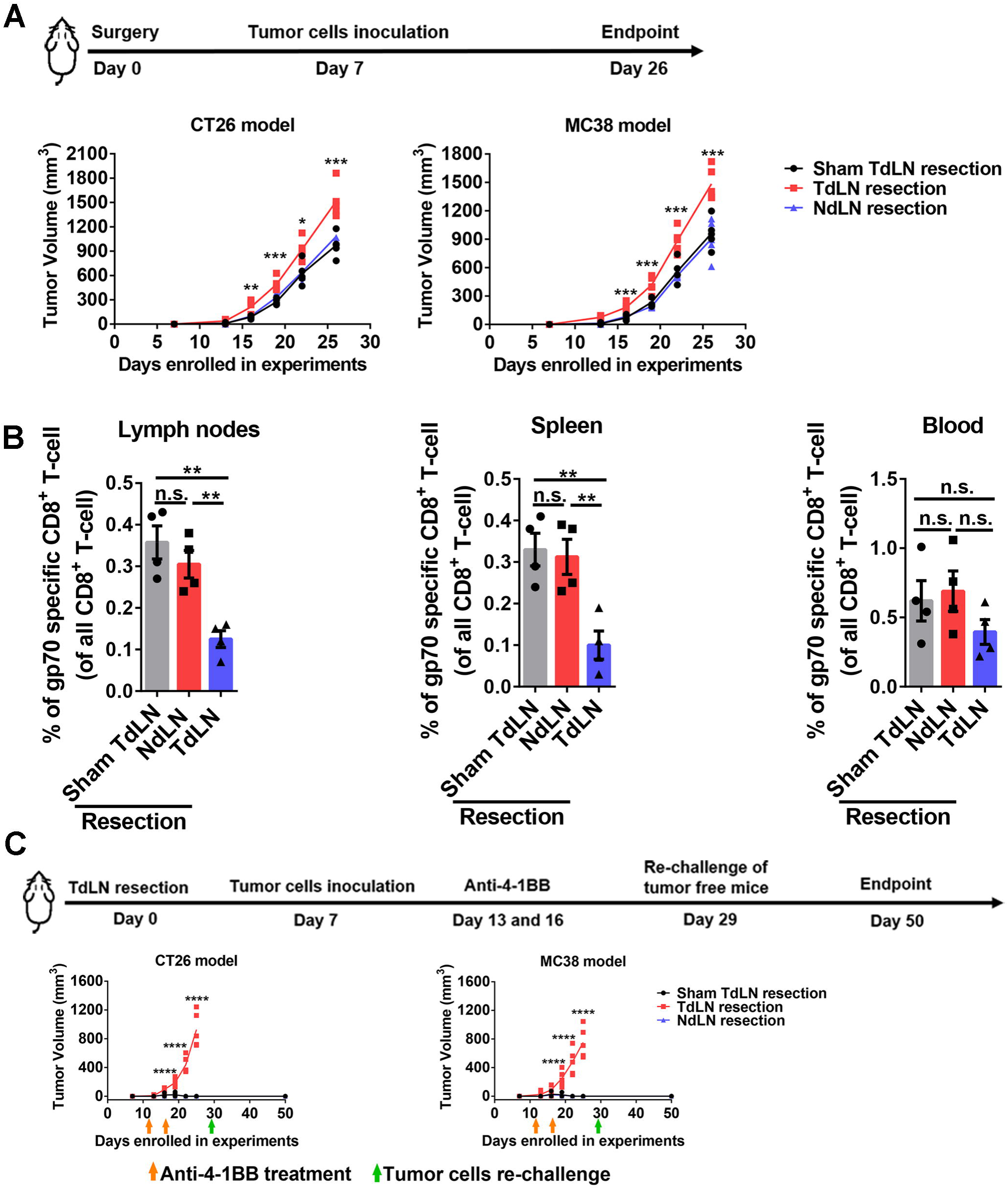
Impact of TdLNs on tumor initiation and immunotherapy response in early-stage tumor models. **A)** Experimental schedule and tumor growth curves in mice with or without TdLNs. Both CT26 (BALB/c mouse as the host) and MC38 (C57BL/6 mouse as the host) subcutaneous models were enrolled in the experiment. Mice were pre-conditioned by TdLNs resection (right inguinal and axillary LNs), NdLNs resection (left inguinal and axillary LNs), or sham surgery prior to tumor inoculation. Accelerated tumor growth was observed in mice without TdLNs (n=5 in each group, One-way ANOVA test between all groups, data represent each individual mouse, \**p*<0.05, \*\**p*<0.01, \*\*\**p*<0.001). **B)** Distribution of tumor antigen (gp70) specific CD8^+^ T cells in tumor-bearing mice with or without TdLNs. Less tumor antigen-specific CD8^+^ T cells was detected in right and left brachial lymph nodes and spleen of TdLNs resected tumor-bearing mice (n=4 in each group, Tukey’s multiple comparisons test between two groups, data were displayed as means ± SEMs, n.s.: no significance, \*\**p*<0.01). **C)** Experimental schedule and early-stage tumor response to anti-4-1BB treatment. Two injections of anti-4-1BB were given shortly after tumor inoculation. The treatment prevented tumor development in tumor-bearing mice with intact TdLNs. Rechallenge of the tumor cells did not form tumors in all anti-4-1BB cured mice (n=4 in each group, One-way ANOVA test between all groups, data represent each individual mouse, \*\*\*\**p*<0.0001). See also Figure S1-S3 ad Table S1.

### TdLNs are not necessary for immunotherapy response in advanced disease tumor models

Since recurrence after primary tumor resection is one of the major causes for treatment failure, we evaluated the impact of TdLNs on tumor recurrence and the response to immunotherapy in mouse models after advanced primary tumor resection. We allowed the primary tumor to grow to a relatively large volume and then resected the primary tumor with and without concurrent TdLN resection. Secondary tumors were then inoculated to mimic localized tumor recurrence (Figure 2A). We confirmed a clean primary tumor resection margin in our models (Table S1), allowing all secondary tumors to start with a comparable baseline. TdLNs were also subjected to histological analysis to confirm that no metastasis developed in TdLNs (Figure S2K). Notably, in our well-controlled model, the secondary tumor growth rate was similar in mice with and without TdLNs (Figure 2A, B). In another group of TdLNs resected mice, we depleted T cells to study the impact of systemic immunity on subsequent tumor development. As predicted, the secondary tumor developed rapidly in mice with impaired systemic immunity (Figure 2A, B). Together, these results indicate that tumor recurrence is accelerated by impaired systemic immunity but not by impaired regional immunity (TdLNs resection).

**Figure 2.**
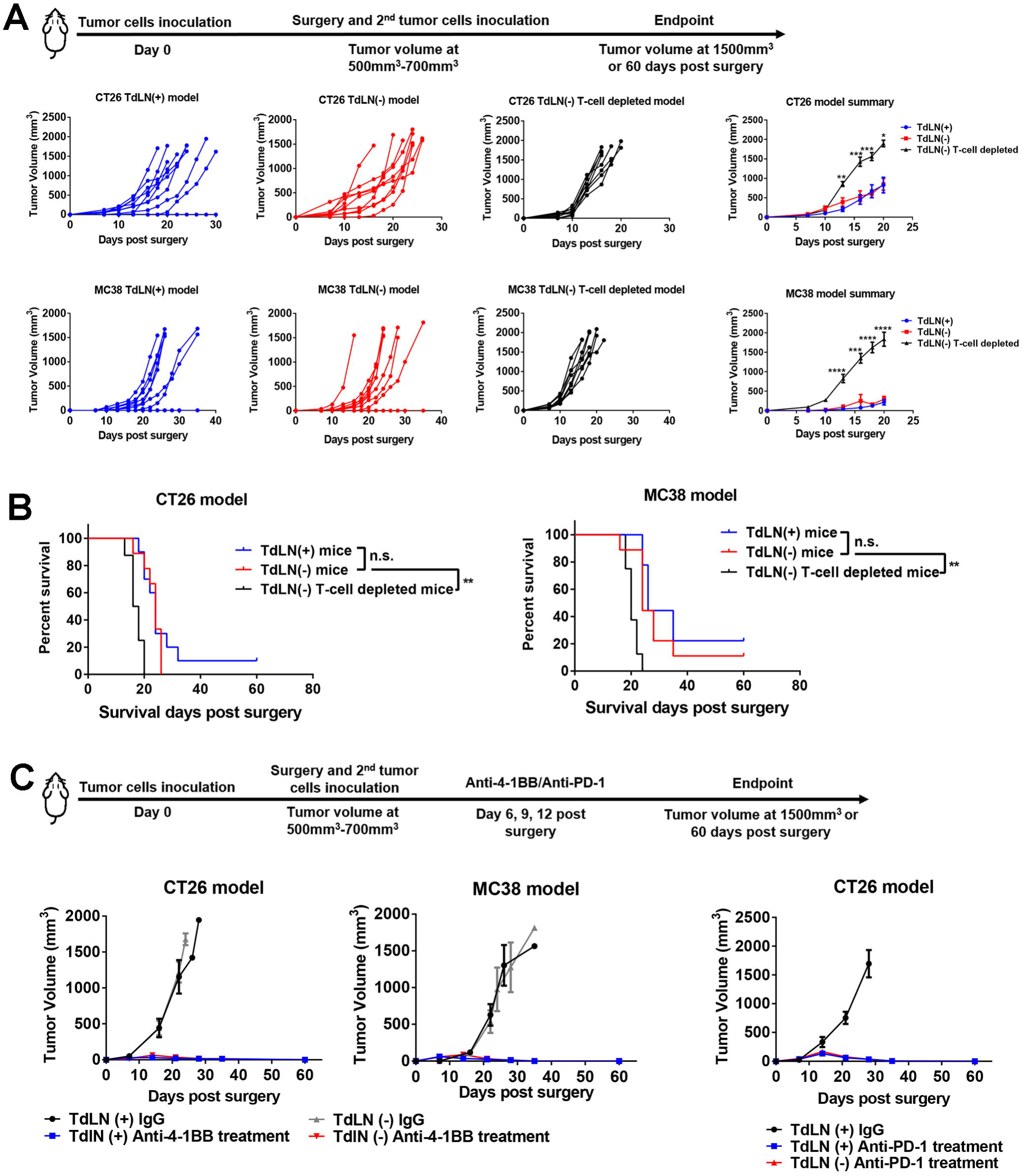
Impact of TdLNs on tumor recurrence and immunotherapy response in advanced stage tumor models. **A)** The experimental schedule. Resection of TdLNs did not accelerate localized secondary tumor (mimicking recurrent tumor) development in both CT26 and MC38 subcutaneous tumor models. However, systemic deletion of T cells significantly accelerated secondary tumor development in both tumor models (n = 8-10 in each group, both individual and summarized curves were shown, t-tests were performed between the TdLN resected and T-cell depleted groups, data were displayed as means ± SEMs, t-test was performed between the TdLN(-) and TdLN(-) T-cell depleted groups, \**p*<0.05, \*\**p*<0.01, \*\*\**p*<0.001, \*\*\*\**p*<0.0001). **B)** Systemic depletion of T cells, but not TdLN resection, led to a shorter survival time of mice due to secondary tumor development (n= 8-10 in each group, log-rank test between indicated groups, \*\**p*<0.01). **C)** Response to anti-4-1BB and anti-PD-1 treatment was tested in localized secondary tumors with or without TdLNs. Anti-4-1BB and anti-PD-1 treatments suppressed secondary tumor growth in both TdLN intact and resected mice (n=5 in each group, data were displayed as means ± SEMs). See also Figure S1, S2, S4, and Table S1.

Next, we asked whether TdLNs resection altered immune infiltration in secondary tumors. The major immune cell types were evaluated in secondary tumors (Figure S4). Total tumor-infiltrating T cells, PD-1 high expression T cells, and MDSCs were not altered in secondary tumors either with or without TdLNs. PD-L1 expression was similar. The frequency of CD103^+^ DCs and lymphatic endothelial cells were significantly higher in the secondary MC38 tumors with TdLNs (Figure S4). However, in the CT26 model, only the lymphatic endothelial cell frequency was statistically higher in secondary tumors with TdLNs than without TdLNs. The frequency of CD103^+^ DCs showed a similar trend but did not reach statistical significance (Figure S4).

Immunotherapies are typically prescribed to patients who have undergone advanced primary tumor resection. In another pre-clinical model, we administrated anti-4-1BB and anti-PD-1 to study whether TdLN resection will lead to immunotherapy resistance. To mimic clinical conditions, we resected the established primary tumor both with and without concurrent TdLN resection. We then inoculated the secondary tumor to mimic localized tumor recurrence. A 6-day gap was allowed between the secondary tumor inoculation and any treatment (Figure 2C). This allows the tumor to connect with systemic circulation and to establish the tumor microenvironment. Then, the mice were treated with anti-4-1BB or anti-PD-1 (Figure 2C). Notably, both anti-4-1BB and anti-PD-1 treatments were efficient in controlling secondary tumor initiation. Secondary tumor control was maintained after TdLN resection (Figure 2C), suggesting that TdLN resection may not be a major influencing factor on the efficacy of ICBT when used as adjuvant therapy in late-stage disease.

### TdLNs shift from an immunoactive to an immunotolerant environment and tumor-antigen specific T cells disseminate during tumor development

Based on the above results, we then hypothesized that immunosuppression in TdLNs and systemic spreading of tumor antigen-specific T cells during tumor development make the TdLNs less important for late-stage tumors compared with early-stage tumors. We collected the TdLNs and NdLNs at different stages of tumor development for analysis and compared them with the naïve LNs. The frequency of CD62L^-^ CD4^+^ T cells was significantly higher in TdLNs than in NdLNs when tumors were small. However, the differences disappeared once the tumors became large (Figure 3A). CD80, a crucial co-stimulatory molecule was higher on APCs in TdLNs than in NdLNs and naïve LNs at early-stage disease (Figure 3B). However, with tumor development, the CD80 level on APCs in TdLNs dropped (Figure 3B). As the receptor of CD80, CD28 is highly expressed on CD4^+^ and CD8^+^ T cells in TdLNs of early-stage tumors but decreased dramatically during tumor development (Figure 3C). Previous studies showed that CD28 is downregulated in T cells which are repetitively exposed to antigens (Lake et al., 1993; Vallejo et al., 1999). Therefore, high numbers of T cells with lower CD28 levels may be the product of repeated activation in the TdLNs of late-stage tumors. However, recent studies indicated that sustained CD28 expression after T-cell priming is required for T-cell function and response to further stimulations, including immune checkpoint inhibitors (Kamphorst et al., 2017; Linterman et al., 2014). IFN-γ is highly produced by functional T cells. However, decreased IFN-γ concentration was observed in TdLNs during tumor development (Figure 3D). These data suggested that immune cells in TdLNs of late-stage tumors may not function as properly as in the TdLNs of early-stage tumors, shifting the TdLNs from an immunoactive to the immunotolerant environment during tumor development.

**Figure 3.**
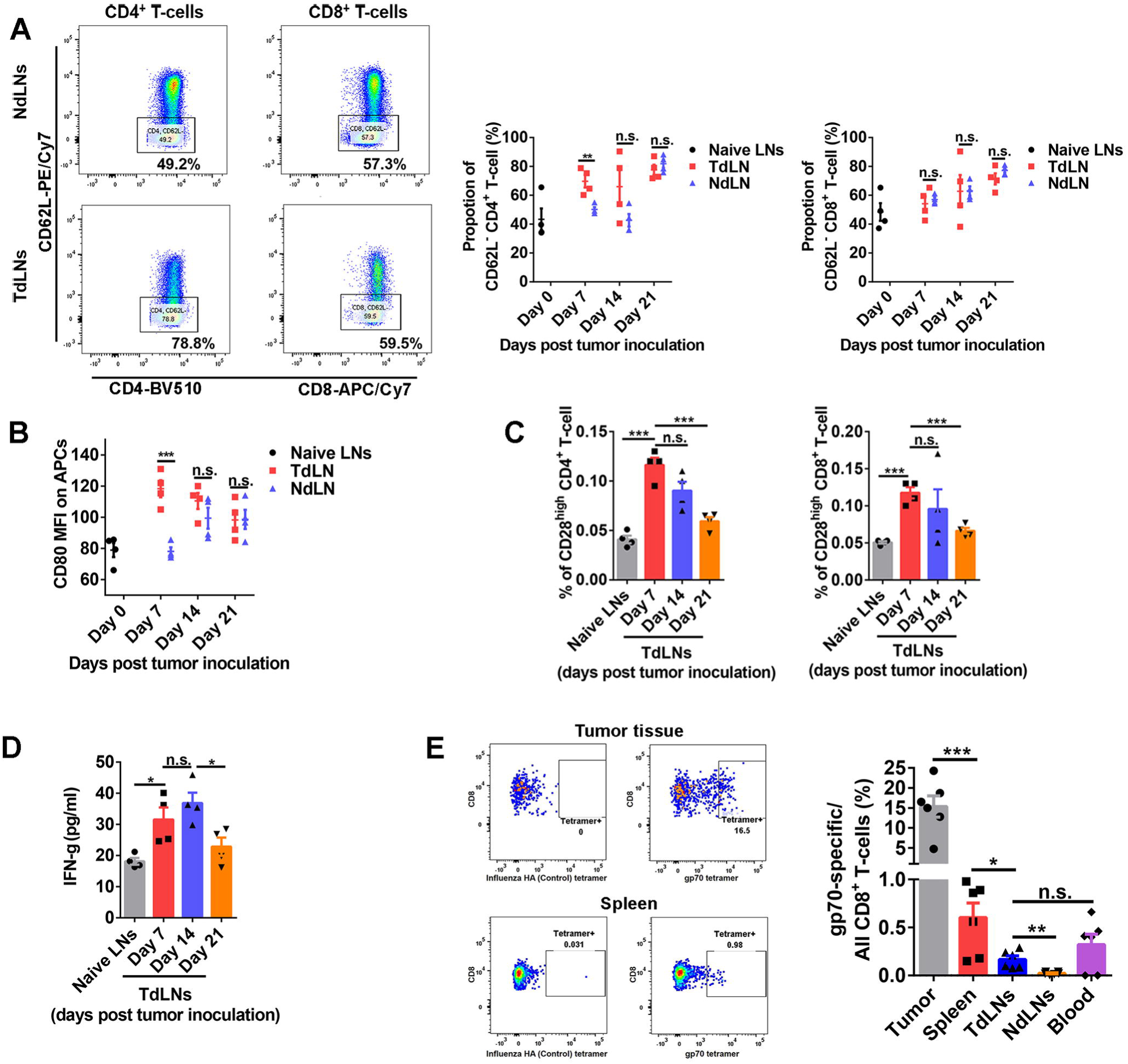
Functional status of TdLNs and tumor antigen-specific T-cell distribution in tumor-bearing mice with advanced disease. **A)** More activated (CD62L^-^) CD4^+^ T cells were observed in TdLNs than NdLNs on 7 days post tumor cells inoculation. However, at the late stage of tumor development, the proportion of activated CD4^+^ T cells was similar in TdLNs and NdLNs. The proportion of activated CD8^+^ T cells were close in TdLNs and NdLNs during tumor development (n=4 in each group, t-tests were performed, data were displayed as means ± SEMs, n.s.: no significance, \*\**p*<0.01). **B)** CD80 expression level on antigen presentation cells (APCs) was higher in TdLNs than NdLNs on 7 days post tumor cells inoculation (n=4 in each group, t-tests were performed, data were displayed as means ± SEMs, n.s.: no significance, \*\*\**p*<0.001, MFI: mean fluorescent intensity). **C)** The proportion of CD28^high^ T cells (both CD4^+^ and CD8^+^) in TdLNs was decreased during tumor development (n=4 in each group, t-tests were performed, data were displayed as means ± SEMs, n.s.: no significance, \*\*\**p*<0.001). **D)** The concentration of IFNγ in TdLNs was higher at the early stage of tumor development than the late-stage (n=4 in each group, t-tests were performed, data were displayed as means ± SEMs, n.s.: no significance, \**p*<0.05). **E)** At the established tumor model (volume 500-700mm^3^), systemic distribution of tumor antigen (gp70) specific CD8^+^ T cells were detected in multiple lymphatic organs and the tumor microenvironment. The tumor microenvironment has the highest frequency of gp70 specific CD8^+^ T cells than lymphatic organs (n=6 in each group, t-tests were performed between indicated groups, data were displayed as means ± SEMs, n.s.: no significance, \**p*<0.05, \*\**p*<0.01, \*\*\**p*<0.001). See also Figure S1-S3 and Table S1.

The amount and distribution of tumor antigen-specific T cells also influence antitumor immunity and immunotherapy response (Liu et al., 2016). We measured the distribution of tumor antigen-specific T cells in mice with established tumors (volume 500-700mm^3^) (Figure S3). As expected, the frequency of gp70 specific CD8^+^ T cells was highest in the tumor-infiltrating CD8^+^ T cells population. The gp70 specific CD8^+^ T cells were detected in all major peripheral lymphatic organs, including spleen, TdLNs, NdLNs, and blood (Figure 3E). The proportion of CD4^+^ (around 55%-65% of all T cells) and CD8^+^ (around 25%-35% of all T cells) T cells in TdLNs was comparable with that in the naïve mice LNs (data not shown). Considering that TdLNs are only a very small proportion of the lymphatic system, our data suggested that in advanced tumor conditions, TdLNs are not the primary reservoir of tumor antigen-specific T cells. The widely distributed tumor antigen-specific T cells in peripheral lymphatic organs could be the responders of immunotherapies for controlling localized residual tumor (minimal secondary tumors in our model) recurrence.

### Sequential treatment of 5-FU and anti-4-1BB or anti-PD-1 leads to better responses than concurrent treatment

In addition to the primary tumor and TdLN resection, chemotherapy is a critical factor potentially affecting the efficacy of immunotherapies. Since our preceding data showed that TdLN resection may not affect the immunotherapeutic response, we then focused on the impacts of chemotherapy on immunotherapies. Several mechanisms by which chemotherapies regulate anti-tumor immunity have been identified (Emens and Middleton, 2015; Fend et al., 2017; Galluzzi et al., 2017; Pfirschke et al., 2016). However, no study has analyzed whether the schedule of combining chemotherapies with immunotherapies influences their synergetic effects. To investigate the impact of different combination therapy schedules on tumor response, we compared sequential versus concurrent 5-FU and anti-4-1BB or anti-PD-1 therapy in mouse models. The IgG and anti-4-1BB monotherapy in immunocompetent and T-cell depleted mice served as control groups (Figure 4A). In mice with established tumors, anti-4-1BB monotherapy delayed tumor growth and prolonged mice survival time (Figure 4B, C). Anti-CD3 impaired systemic immunity by suppressing T-cell populations (Figure S5). In an established tumor model, anti-CD3 preconditioning nullified the anti-tumor effects of anti-4-1BB (Figure 4B, C), indicating that intact systemic immunity was required for anti-4-1BB response. 5-FU also delayed tumor development in established tumor models (Figure 4B, C). We then combined anti-4-1BB with the 5-FU treatment and found no noticeable improvement in mice survival time (Figure 4B, C). In another cohort of mice, the 5-FU treatment was used as induction, and then later, anti-4-1BB was added as the maintenance treatment (Figure 4B, C). To determine an appropriate sequential treatment strategy, we tested the dynamics of 5-FU induced T-cell depletion (Figure S5). In the sequential treatment, anti-4-1BB was given when the T-cell population had almost recovered from the 5-FU treatment. Mice treated with sequential combination therapy had the longest survival time and the most effective tumor control of all cohorts (Figure 4B, C).

**Figure 4.**
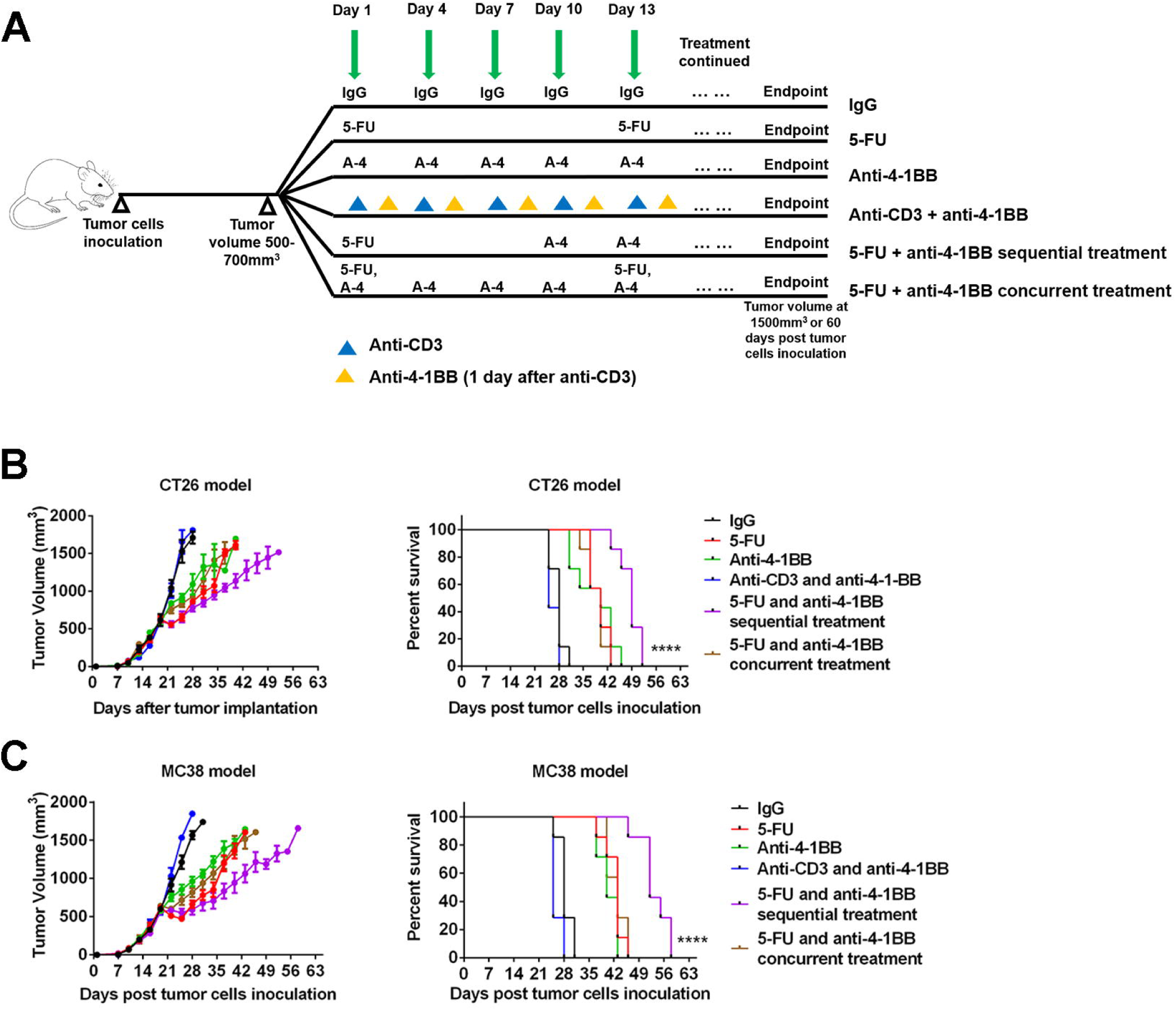
5-FU and anti-4-1BB sequential treatment elicits strong antitumor activity. **A)** Tumor (500mm^3^ to 700mm^3^ in volume) bearing mice were randomly assigned to 6 treatment groups: IgG (one dose/3 days), 5-FU monotherapy (one dose/12 days), anti-4-1BB monotherapy (one dose/3 days), anti-CD3 therapy (one dose/3 days) and anti-4-1BB therapy (one dose/3 days, two days after anti-CD3), 5-FU (one dose) and anti-4-1BB (1 dose/3 day, starting at 9 days post 5-FU) sequential therapy, and 5-FU (one dose/12 days) and anti-4-1BB (1 dose/3 day, started at the same day of 5-FU) concurrent therapy. The treatment was continued until the endpoint of follow-up. **B)** CT26 tumor response to different treatments. The 5-FU and anti-4-1BB sequential treatment significantly prolonged survival time of the tumor-bearing mice (n=4 in each group for the tumor growth curve, data were displayed as means ± SEMs, n=7 in each group for the mouse survival curve, log-rank test between all survival curves, \*\*\*\**p*<0.0001). **C)** The same experiments of panel B were repeated in the MC38 tumor model (n=4 in each group for the tumor growth curve, data were displayed as means ± SEMs, n=7 in each group for the mouse survival curve, log-rank test between all survival curves, \*\*\*\**p*<0.0001). See also Figure S5 and S6.

Next, we compared the 5-FU and anti-4-1BB sequential and concurrent treatments in a more clinically relevant model. In this model, we performed resection of the established primary tumor together with its TdLNs and induced localized secondary tumors for treatment (Figure 5A). Over 60 days of the experiment, the sequential treatment showed better tumor suppression than concurrent treatment (Figure 5B). Anti-PD-1 is an FDA-approved class of cancer-directed immunotherapy with different mechanisms than anti-4-1BB. To test whether the conclusion from the anti-4-1BB treatment was generalizable to the anti-PD-1 treatment, we combined 5-FU and anti-PD-1 in concurrent and sequential schedules. Again, the 5-FU and anti-PD-1 given in sequence showed better tumor control than when administered concurrently (Figure 5C).

**Figure 5.**
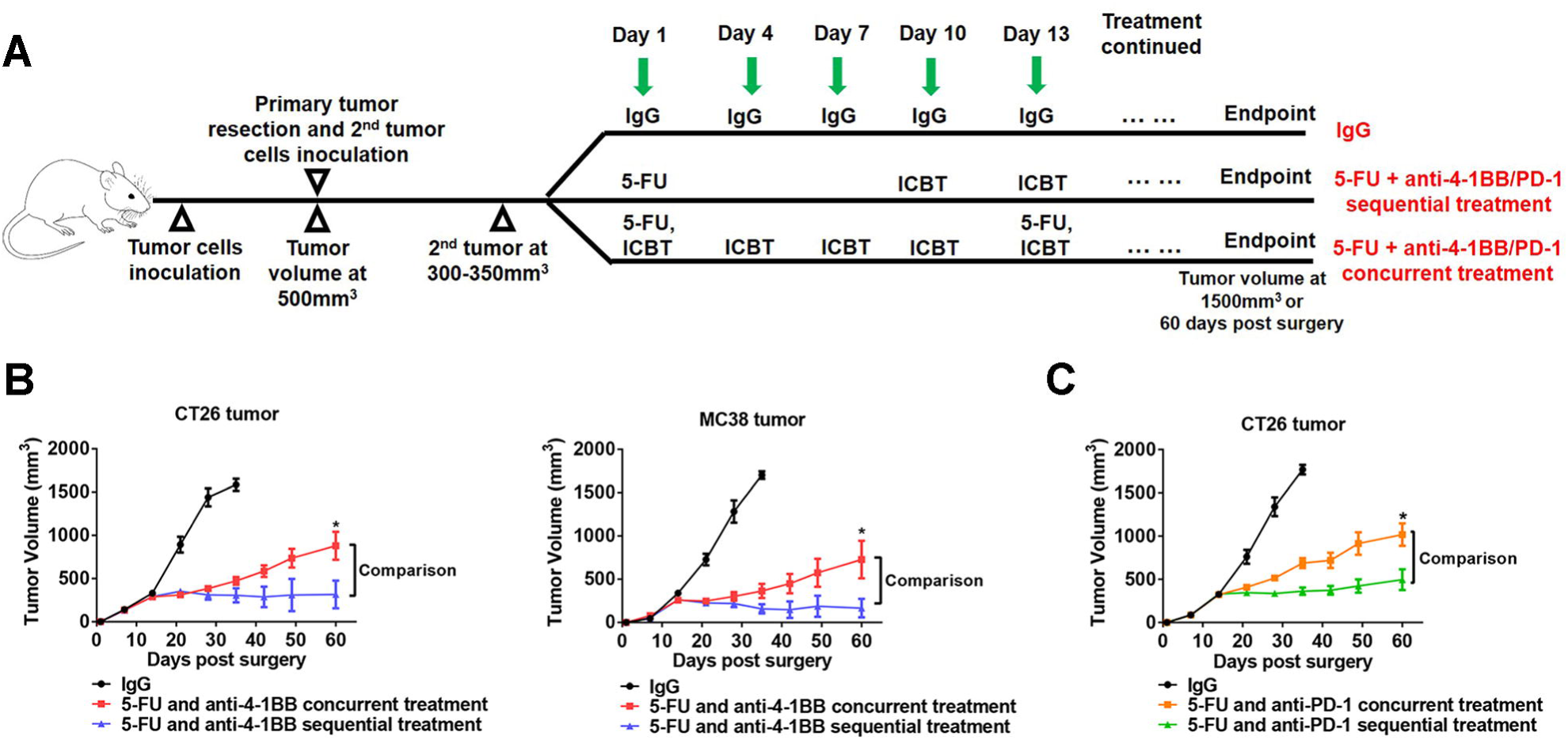
5-FU and anti-4-1BB sequential treatment on secondary tumors that mimic tumor recurrence. **A)** The primary tumor and TdLNs were resected when tumors are at around 500mm^3^ in volume. Secondary tumors were induced and treated by different strategies at 300mm^3^-350mm^3^ in volume. Some mice rejected the secondary tumors and were not included in the therapeutic study. **B-C)** The 5-FU and anti-4-1BB or anti-PD-1 sequential treatment was more efficient than the 5-FU and anti-4-1BB or anti-PD-1 concurrent treatment in controlling secondary tumors in CT26 and MC38 models (n=7 in each group, data were displayed as means ± SEMs, t-test was performed between the sequential and concurrent treatment groups, * *p*<0.05, ICBT: immune checkpoint blockade therapy (anti-4-1BB or anti-PD-1)).

Toxicity is a primary concern for cancer treatments, especially in combination therapy. We took this into account by evaluating the side effects of each treatment. 5-FU monotherapy and 5-FU and anti-4-1BB concurrent combination therapy caused severe body weight loss and diarrhea during the treatment (Figure S6). In contrast, the 5-FU and anti-4-1BB sequential combination therapy showed slight or no side effects for the duration of the experiment (Figure S6).

### Sequential treatment of 5-FU and anti-4-1BB or anti-PD-1 stimulates a strong antitumor immune response

Our pre-clinical models suggested that 5-FU and anti-4-1BB or anti-PD-1 sequential treatment has superior tumor controlling effects than the concurrent treatment schedule. We investigated the potential mechanisms of this result by performing mass cytometry to generate a comprehensive immune landscape characterization in tumor tissues (Figure S7). Notably, CD80 and CD86 expression were upregulated after 5-FU and anti-4-1BB sequential treatment in CT26 tumors (Figure 6A). High expression of these two critical co-stimulatory factors suggests enhanced tumor visibility by T cells. The expression of PD-L1 on tumor tissue was not significantly changed among different groups (Figure 6A). Furthermore, tumor immune infiltration studies showed that anti-4-1BB monotherapy stimulated tumor-infiltrating T-cell proliferation and increased the CD8^+^ T-cell versus regulatory T-cell ratio (Figure 6B, C, D). In sum, these experiments showed that 5-FU and anti-4-1BB sequential treatment alone maintained the positive effects of anti-4-1BB on the T cell population (Figure 6B, C, D).

**Figure 6.**
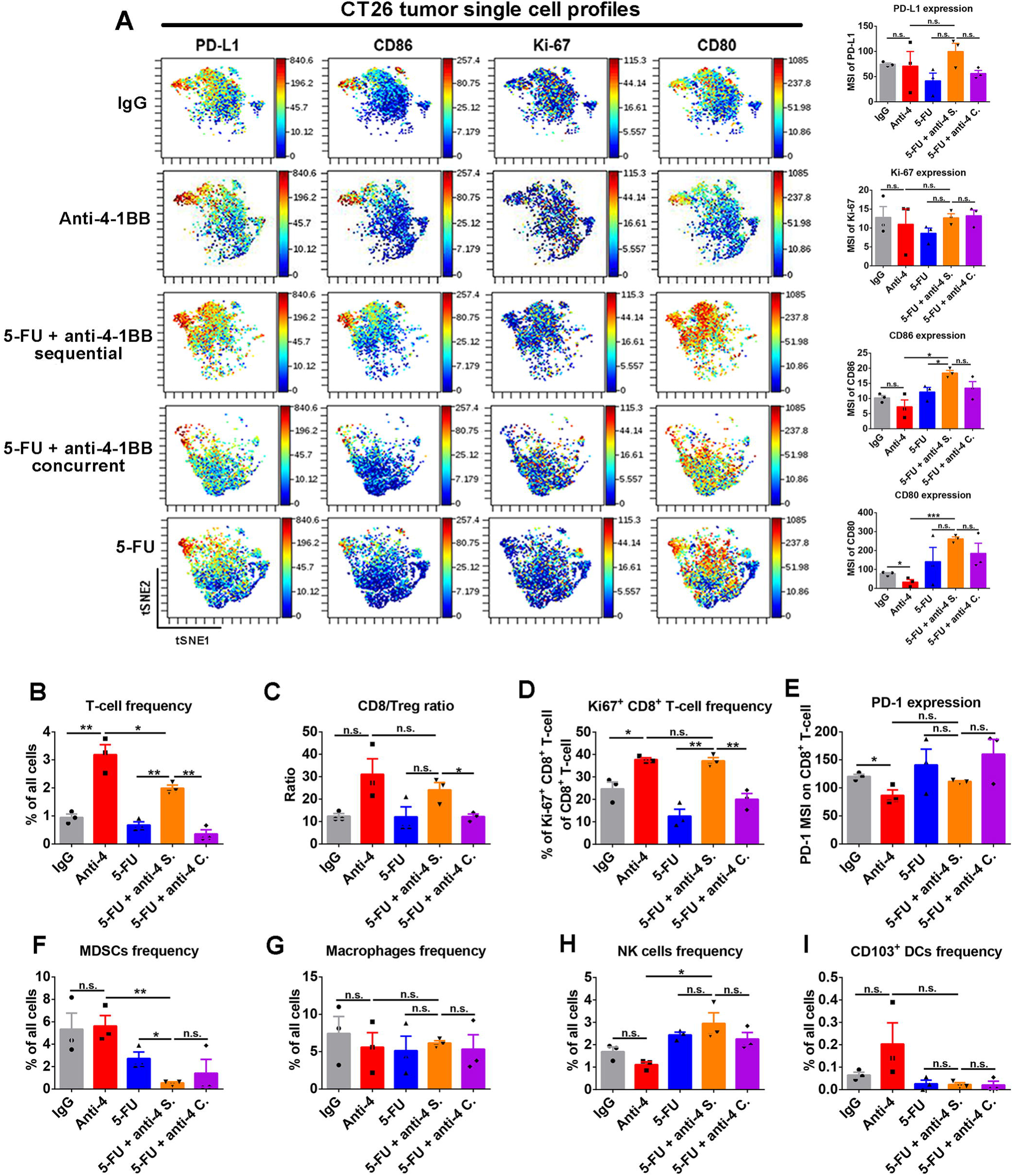
Tumor immunological response to 5-FU and anti-4-1BB treatments in CT26 tumors. **A)** ViSNE plot showed single cell level expression of PD-L1, Ki-67, CD80, and CD86 in the tumor tissue. PD-L1, Ki-67, CD80, and CD86 expression were quantified in whole tumor tissue. The 5-FU and anti-4-1BB sequential treatment significantly upregulated CD86 and CD80 expression in tumor tissues (n=3 in each group, data were displayed as means ± SEMs, t-test was performed between the indicated groups, n.s.: no significance, \**p*<0.05, \*\*\**p*<0.001, MSI: mean signal intensity). **B-I)** The tumor-infiltrating T cells frequency, CD8/Treg ratio, Ki-67^+^ CD8^+^ T cells frequency, expression of PD-1 on CD8^+^ T cells, myeloid-derived suppressive cells (MDSCs) frequency, macrophages frequency, NK cells frequency, and CD103+ dendritic cells (DCs) frequency were measured in tumors treated by different strategies (n=3 in each group, data were displayed as means ± SEMs, t-test was performed between the indicated groups, n.s.: no significance, \**p*<0.05, \*\**p*<0.01, MSI: mean signal intensity). See also Figure S7-9.

Meanwhile, the tumors treated by the sequential therapy had the lowest MDSC frequency and highest NK cell frequency (Figure 6F, H). PD-1 expression on CD8^+^ T cells and macrophages frequency was similar among all groups (Figure 6E, G). CD103^+^ DC frequency trended higher in the anti-4-1BB monotherapy group, but was not statistically significant (Figure 6I). We repeated the same experiment in MC38 tumors (Figure S8) and obtained similar results as in CT26 tumors with most parameters tested. However, CD80 and CD86 expression levels in MC38 tumors were not increased significantly by 5-FU and anti-4-1BB sequential treatments. This difference between MC38 and CT26 tumors indicates the tumor-dependent effects of the treatment. In CT26 tumors, we also evaluated the immunoregulatory effects of 5-FU and anti-PD-1 combination (Figure S9). Tumor-wide expression of PD-L1, CD86, and CD80 was increased in 5-FU and anti-PD-1 sequential treatment group. In addition, the frequencies of total T cells, proliferating CD8^+^ T cells and NK cell was highest in tumors treated by 5-FU and anti-PD-1 sequential therapy (Figure S9). Notably, the frequency of MDSCs was decreased by 5-FU monotherapy and combination treatment (Figure S9). These findings showed the immunological impacts of different treatment strategies and reinforced that using 5-FU as an induction treatment and then anti-4-1BB or anti-PD-1 as maintenance treatments produces the most prominent and synergic effect in reversing the immunosuppressive tumor microenvironment.

## DISCUSSION

Immunotherapies are mostly used as second- or third-line treatments for treatment-refractory tumors. However, studies that investigate the impact of different clinical conditions and combination strategies on tumor immunotherapy are limited. Here, we comprehensively profiled the impacts of tumor-draining lymph nodes (TdLNs) resection and different chemotherapy combination schedules on ICBT responses.

Surgery has been a dominant strategy for several decades to prevent, diagnose, stage, and treat cancers. Radical surgery—a procedure that removes blood supply to the tumor, lymph nodes and sometimes adjacent structures—is routinely performed in many cancers such as colorectal cancer, breast cancer, and lung cancer. Early-stage cancer patients have excellent disease control with surgery alone, yet advanced diseases require more comprehensive treatments, including chemotherapies, oncogenic pathway targeted therapies, and immunotherapies. Currently, most immunotherapies are used as adjuvant treatments (given after surgeries). TdLNs are the primary lymphatic organs where antitumor immune responses are initiated (Fisher and Fisher, 1971; Jeanbart et al., 2014; Marzo et al., 1999; Munn and Mellor, 2006; Shu et al., 2006). In mouse models with resected TdLNs before tumor cell inoculation, we observed that removal of TdLNs significantly accelerated tumor growth and compromised response to immunotherapy. These data uncovered a key role for TdLNs in preventing cancer cells from evading antitumor immunity at early stages. Mechanistically, TdLNs resection in early-stage disease led to inadequate antitumor immune simulation, featured by a low frequency of tumor antigen-specific T cells in lymphatic organs. Our observations were in line with previous studies, highlighting the significance of TdLNs in initiating antitumor immunity and regulating immunotherapy response in early-stage disease (Fransen et al., 2018).

Recently, Fransen et al reported that TdLNs are determining factors of PD-1/PD-L1 immune checkpoint therapies in early-stage tumor models (Fransen et al., 2018). However, whether the TdLNs are critical for immunotherapy response in recurrent tumor models, which represent a major clinical issue, had not been addressed. In our study, we established a model to mimic tumor recurrence from residual tumor lesions after primary tumor and TdLN resection. We first thoroughly resected the primary tumors either with or without TdLN resection and confirmed a clean surgical margin. We then inoculated tumor cells *in situ* to induce a secondary tumor. This method allows all secondary tumors to have a relatively similar baseline volume and growth dynamic before any treatment. We also allow the localized secondary tumors to connect with systemic circulation and establish a tumor microenvironment before treatment was initiated. Our well-designed model provided a platform for an unbiased evaluation of treatment efficacy in residual disease after primary tumor resection.

With our model, we found that resection of TdLNs in advanced tumors did not influence localized secondary tumor immunity and response to immunotherapies (anti-PD-1 and anti-4-1BB). Furthermore, we investigated the factors that determine the significance of TdLNs in antitumor immunity and immunotherapeutic response. Previous findings indicated that the bidirectional crosstalk between tumor cells and TdLNs allowed remodeling of each other during tumor progression (Fisher and Fisher, 1971; Ito et al., 2006; Munn and Mellor, 2006; Shu et al., 2006; Watanabe et al., 2008). Immunosuppressive factors derived from tumors such as TGFβ, can drain to TdLNs and induce an immunosuppressive microenvironment (Cochran et al., 2006; Ito et al., 2006). We tested the hypothesize that antitumor function of TdLNs is impaired in advanced tumor models. We compared the immune responses in naïve LNs, TdLNs of early-stage and advanced tumors and demonstrated a trend between potent immunosuppression in TdLNs and tumor progression. Although the TdLNs eventually became immunotolerant, the distribution of tumor antigen-specific T cells are extensive in lymphatic tissues in advanced tumors. Resection of TdLNs did not significantly reduce the population of tumor-antigen specific T cells that respond to immunotherapies. Our data corroborate with previous reports showing strong immunosuppression development in TdLNs of human cancers (Murthy et al., 2019; Shuang et al., 2017). This explains why resection of TdLNs may not influence the antitumor immunity in late-stage tumor models. Finally, it is also important to understand that the resected TdLNs in our experimental models might have developed immunotolerance. However, since humans have more TdLNs than the mouse model, immunoactive TdLNs do exist in certain circumstances and might influence immunotherapy response (Toki et al., 2020; Wu et al., 2014). Therefore, it will be critical to evaluate the functional status of TdLNs in humans before extending our conclusions to human cancers.

Systemic therapies, such as chemotherapies are used to treat primary tumors, eradicate micrometastatic disease, or stabilize the disease in widespread incurable conditions (DeVita and Chu, 2008). Chemotherapies have the advantages of being fast-acting and effective, thus they are widely administered as the primary treatment for combinational strategies (DeVita and Chu, 2008). Combinations of chemotherapies with immunotherapies are extensively discussed and currently tested in pre-clinical models and clinical trials (Emens and Middleton, 2015; Kareva, 2017; Pfirschke et al., 2016; Wang et al., 2018). Comprehensive studies have revealed the mechanisms by which chemotherapy can promote antitumor immunity by induction of immunogenic cell death and disruption of tumor microenvironment components that are used to evade the immune response (Galluzzi et al., 2017; Lutsiak et al., 2005; Michels et al., 2012; Samanta et al., 2018; Tesniere et al., 2010). However, cancer chemotherapies are also considered immunosuppressive due to their cytotoxic effects on immune cells. Therefore, we speculated that the same chemotherapy may have different impacts on anti-tumor immunity, either stimulatory or inhibitory, depending on the specific combination schedules. We used 5-FU, a common chemotherapeutic agent, as a representative agent to study the influences of different chemotherapeutic and immunotherapeutic combination strategies on the anti-tumor immune response.

Through extensive study of 5-FU induced immune responses, we revealed both systemic immunosuppressive effects and immune-stimulating effects in the tumor microenvironment. 5-FU treatment upregulated CD80 expression and depleted MDSCs. CD80 is a protein found on antigen-presenting cells as well as tumor cells and belongs to the B7 family; it provides a costimulatory signal necessary for activating T cells and natural killer cells (Beyranvand Nejad et al., 2016; Chambers et al., 1996; Lanier et al., 1995; Singh et al., 2003). Thus, the upregulation of CD80 in tumor tissue induced by 5-FU treatment will potentially lead to increased tumor visibility by T cells. MDSCs are a heterogeneous population of cells that potently suppress T-cell responses (Kumar et al., 2016; Veglia et al., 2018). By depleting MDSCs in tumor tissue, 5-FU treatment may potentiate antitumor immunity by eliminating the negative regulations. These findings are also supported by a previous report (Vincent et al., 2010). In addition to the immunogenic effects, we also observed that 5-FU treatment suppressed the T-cell population in the tumor microenvironment. Thus, avoiding the immunosuppressive effects and preserving the immunogenic effects of 5-FU treatment will determine the response of 5-FU and immunotherapy combinations.

In our study, the administration of anti-4-1BB or anti-PD-1 after 5-FU treatment significantly improved tumor responses. In this combination strategy, anti-4-1BB or anti-PD-1 selectively boosted response of T-cell and NK cells while the 5-FU treatment increased tumor visibility and suppressed MDSCs. However, when anti-4-1BB or anti-PD-1 was added to the repetitive 5-FU treatment, less synergistic effects were observed. Our data highlighted the importance of determining the best schedule for designing a successful chemo-immunotherapy combination. In addition to timing, dosing is another potential factor that affects the chemotherapy-induced immune response. Low dose chemotherapies have shown special immunoregulatory effects in tumor models (Cao et al., 2014; Ghiringhelli et al., 2007). Further studies are needed to test different chemotherapy doses on the chemo-immunotherapy combination.

In conclusion, our research investigated how traditional cancer treatments will affect novel immunotherapies in clinically relevant tumor models. Our findings indicate that TdLN resection can have adverse impact on anti-tumor immunity, but only in early-stage tumor models. In advanced tumor models, resection of immunotolerant TdLNs during primary tumor surgery does not significantly alter anti-tumor immunity or immunotherapy response in secondary tumors that mimic localized tumor recurrence. Meanwhile, minimizing the immunosuppression and strengthening the immunogenic effects of traditional cancer therapies are critical for immunotherapy induced durable cancer remission. Specifically, sequential cytotoxic chemotherapy followed by immunotherapy produced a significantly higher degree of anti-tumor response compared to concurrent combination therapy. These findings highlight the need to test immunotherapies in tumor models that more closely mimic different clinical conditions and establish references for designing clinical trials to determine the most effective cancer immunotherapy strategies.

### Limitations of the study

Our pre-clinical studies were mainly performed on mouse tumor models. Although mouse models are heavily used in preclinical studies, mouse tumor development, numbers of tumor-draining lymph nodes and disease kinetics are different from human clinical conditions. While we refined our surgical methods and mouse models to closely reflect human conditions, the differences in mouse anatomy and physiology may potentially limit the translational value of our conclusions. Besides, our study was limited only to the commonly used chemotherapies such as 5-FU in pre-clinical models, therefore further clinical trials are needed to validate our findings acquired in pre-clinical settings.

## ACKNOWLEDGMENTS

This work is supported by the Minnesota Colorectal Cancer research Funds, Mezin-Koats Colon Cancer Research Award and the ChainBreaker funds from Masonic Cancer Center and University of Minnesota, Medical School Innovation award and the Department of Surgery, Research funds. We thank the Mass Cytometry core facility at University of Minnesota for helpful assistance. We thank Dr. Matthew Robertson and Ms. Isabella Ramirez for assisting with manuscript editing.

## AUTHOR CONTRIBUTIONS

X.Z. and S.S. conceived and designed the experiments. X.Z., B.K., and D.W. performed all experiments and data analyses. X.Z., B.K., D.W., E.L. and S.S. commented on the data and co-wrote the paper. S.S. supervised this project.

