## Supplemental information for "Chemotherapy but not the tumor draining lymph nodes determine the immunotherapy response in secondary tumors"

### Supplemental Figures and Legends

#### Figure S1

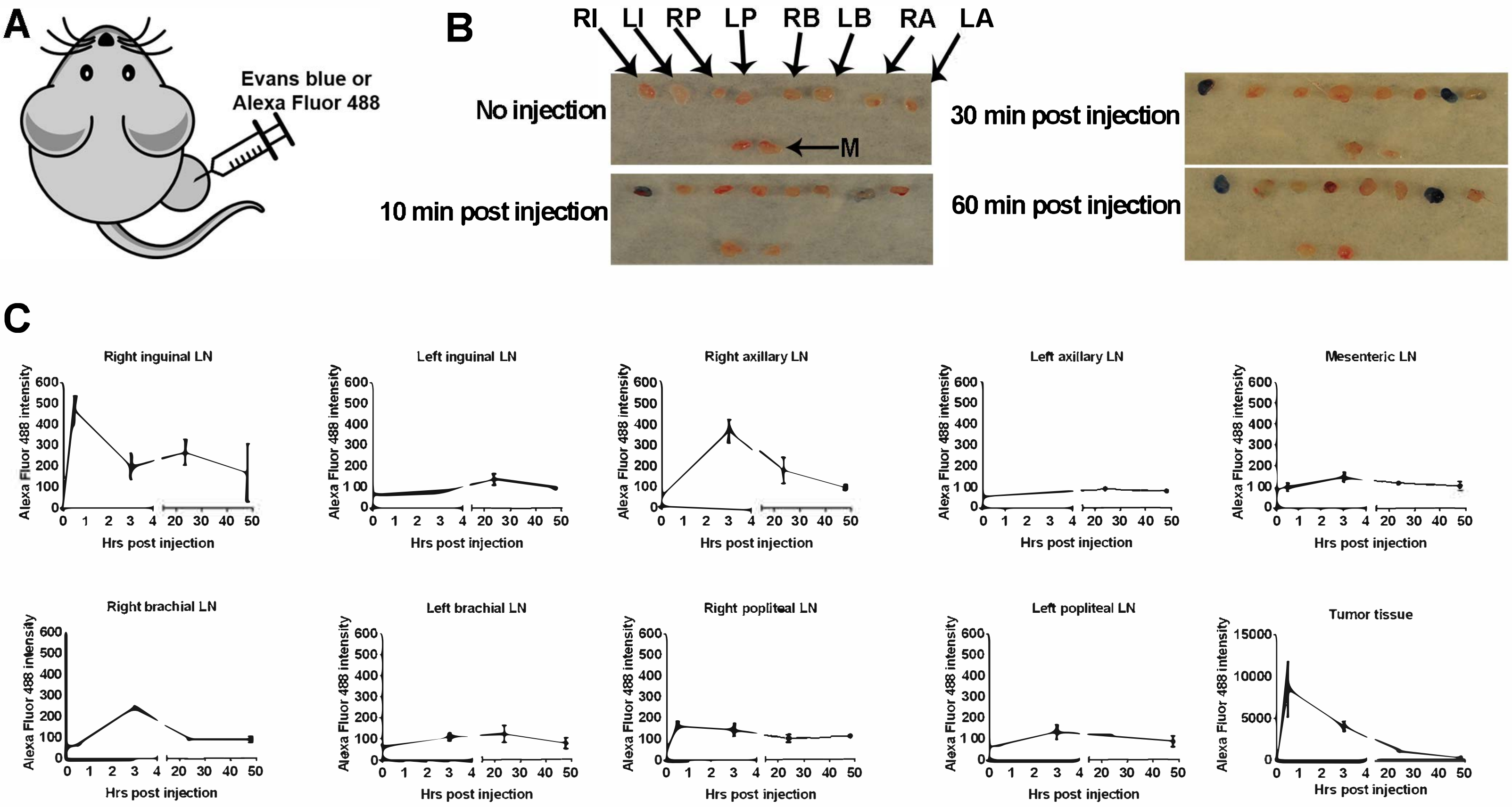

**Figure S1. Identification of tumor-draining lymph nodes (TdLNs) in mouse. Related to Figure 1-3.**
**A)** Evan blue dye or Alexa Fluor® 488 dye was injected into the tumor in the right hinge flank to trace TdLNs.
**B)** 10 min post Evan blue dye injection in the right hinge flank tumor, the right inguinal (RI) and right axillary (RA) lymph nodes (LNs) were stained. The deeper color was seen at 30 min and 60 min post-injection. The representative data from three independent experiments were shown.
**C)** Flow cytometry was used for detecting the Alexa Fluor® 488 dye distribution in lymphatic organs (drainage from the right hinge flank tumor injection site). The RI LN and RA LN showed the highest FITC signal and were identified as the major TdLNs. The other LNs were identified as the non-draining lymph nodes (NdLNs) (n=3 in each group, data displayed as means ± SEMs, RI: right inguinal, LI: left inguinal, RP: right popliteal, LP: left popliteal, RB: right brachial, LB: left brachial, RA: right axillary, LA: left axillary, M: mesenteric).

### Figure S2

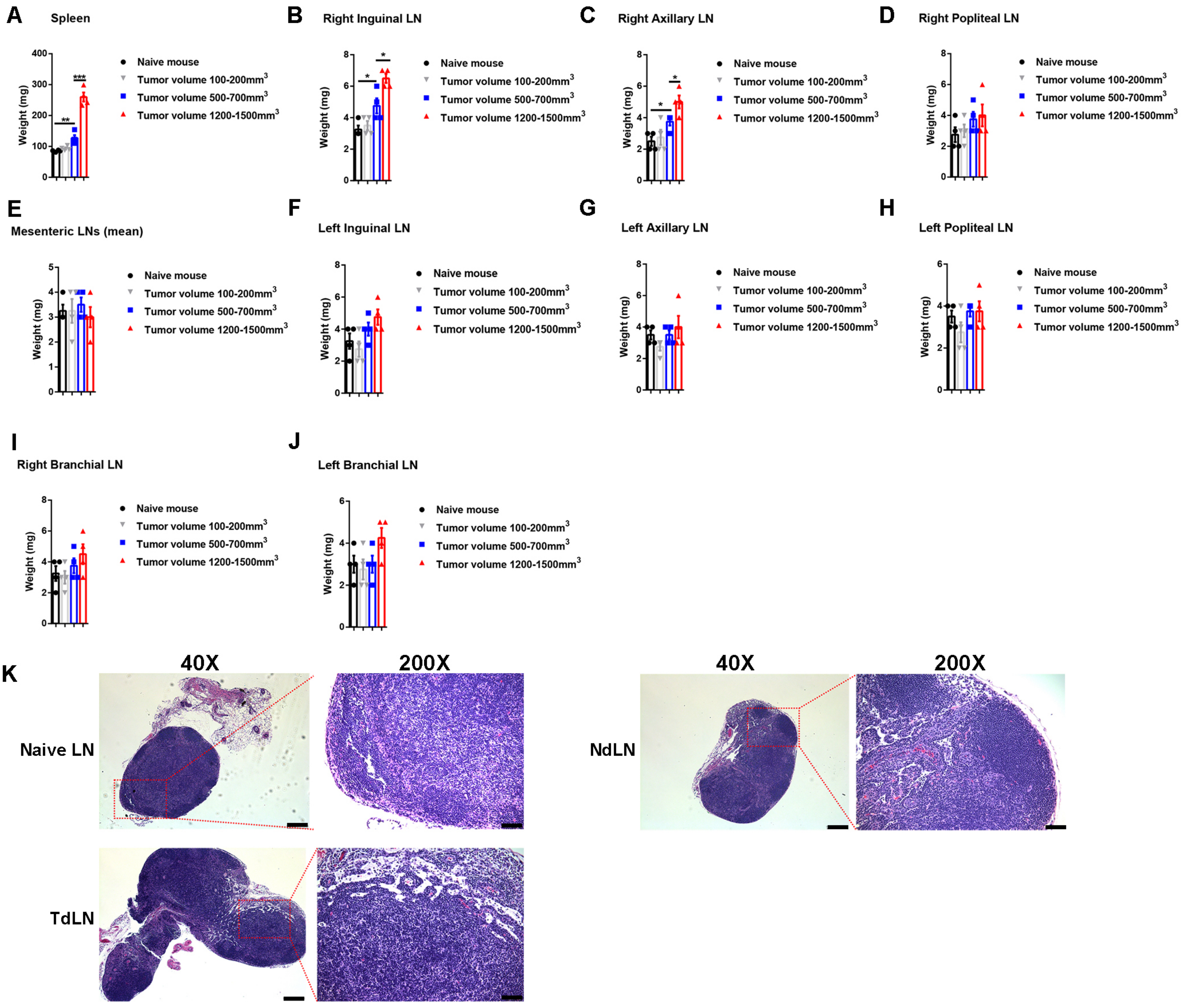
A
Spleen
B
Right Inguinal LN
C
Right Axillary LN
D
Right Popliteal LN
E
Mesenteric LNs (mean)
F
Left Inguinal LN
G
Left Axillary LN
H
Left Popliteal LN
I
Right Branchial LN
J
Left Branchial LN
Weight (mg)
Naive mouse
Tumor volume 100-200mm3
Tumor volume 500-700mm3
Tumor volume 1200-1500mm3
K
40X
200X
Naive LN
TdLN
NdLN

**Figure S2. Physical changes and histology of TdLNs, NdLNs, and spleen of tumor-bearing mice. Related to Figure 1-3.**

**A-J)** The weight of spleen and major superficial LNs (both TdLNs and NdLNs) were measured at different time points of tumor development. During tumor development, a significant splenomegaly was observed. An obvious lymphadenopathy was observed in the TdLNs rather than in the NdLNs during tumor development (n=4 in each group, t-test was performed between indicated groups, data were displayed as means ± SEMs, $^{*}p<0.05$, $^{**}p<0.01$, $^{***}p<0.001$, statistical analyses without significance were not shown).

**K)** At the late-stage of tumor development, the histology of TdLNs and NdLNs was evaluated. The TdLNs were larger than NdLNs and naïve LNs (taken from tumor-free mice). No metastasis was observed in TdLNs. Representative data from three independent experiments were shown (Scale bars: 250μm in 40X images and 50μm in 200X images).

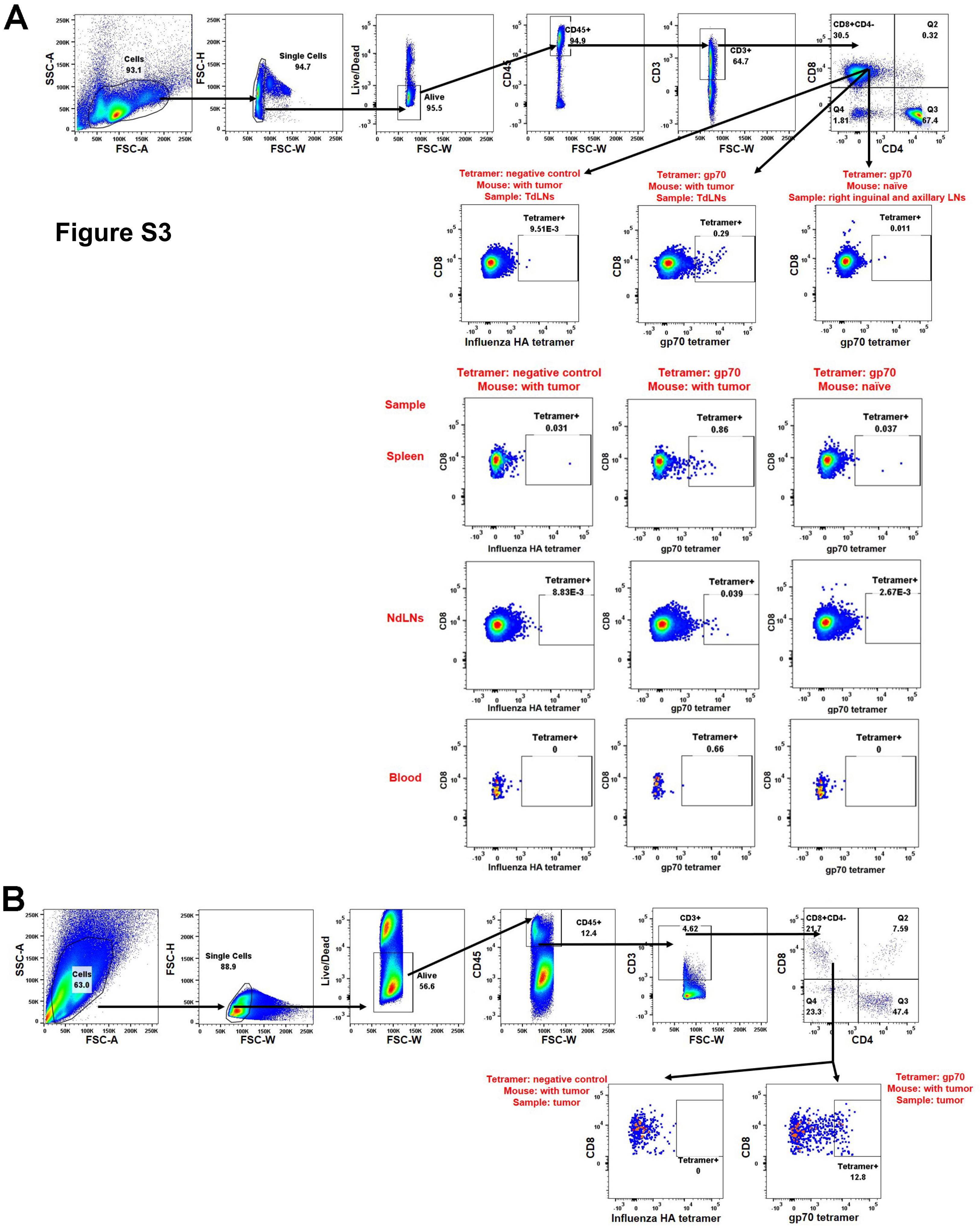
A
SSC-A
FSC-A
Cells
93.1
FSC-H
FSC-W
Single Cells
94.7
Live/Dead
Alive
95.5
CD45
CD45+
94.9
CD3
CD3+
64.7
CD8
CD4
CD8+CD4-
30.5
Q2
0.32
Q4
1.81
Q3
67.4
Tetramer: negative control
Mouse: with tumor
Sample: TdLNs
Tetramer: gp70
Mouse: with tumor
Sample: TdLNs
Tetramer: gp70
Mouse: naïve
Sample: right inguinal and axillary LNs
Tetramer+
9.51E-3
Tetramer+
0.29
Tetramer+
0.011
Influenza HA tetramer
gp70 tetramer
Tetramer: negative control
Mouse: with tumor
Tetramer: gp70
Mouse: with tumor
Tetramer: gp70
Mouse: naïve
Sample
Spleen
Tetramer+
0.031
Tetramer+
0.86
Tetramer+
0.037
NdLNs
Tetramer+
8.83E-3
Tetramer+
0.039
Tetramer+
2.67E-3
Blood
Tetramer+
0
Tetramer+
0.66
Tetramer+
0
B
Cells
63.0
Single Cells
88.9
Alive
56.6
CD45+
12.4
CD3+
4.62
CD8+CD4-
21.7
Q2
7.59
Q4
23.3
Q3
47.4
Tetramer: negative control
Mouse: with tumor
Sample: tumor
Tetramer: gp70
Mouse: with tumor
Sample: tumor
Tetramer+
0
Tetramer+
12.8

**Figure S3**

**Figure S3. Gating of tumor antigen-specific $CD8^+$ T cells. Related to Figure 1 and 3.**
**A)** Representative gating process of $CD8^+$ gp70 (tumor) antigen-specific T cells in peripheral lymphatic organs.
**B)** Representative gating process of $CD8^+$ gp70 (tumor) antigen-specific T cells in tumor tissues. The negative tetramer and lymphatic organs from naïve mice were used as controls for gating.

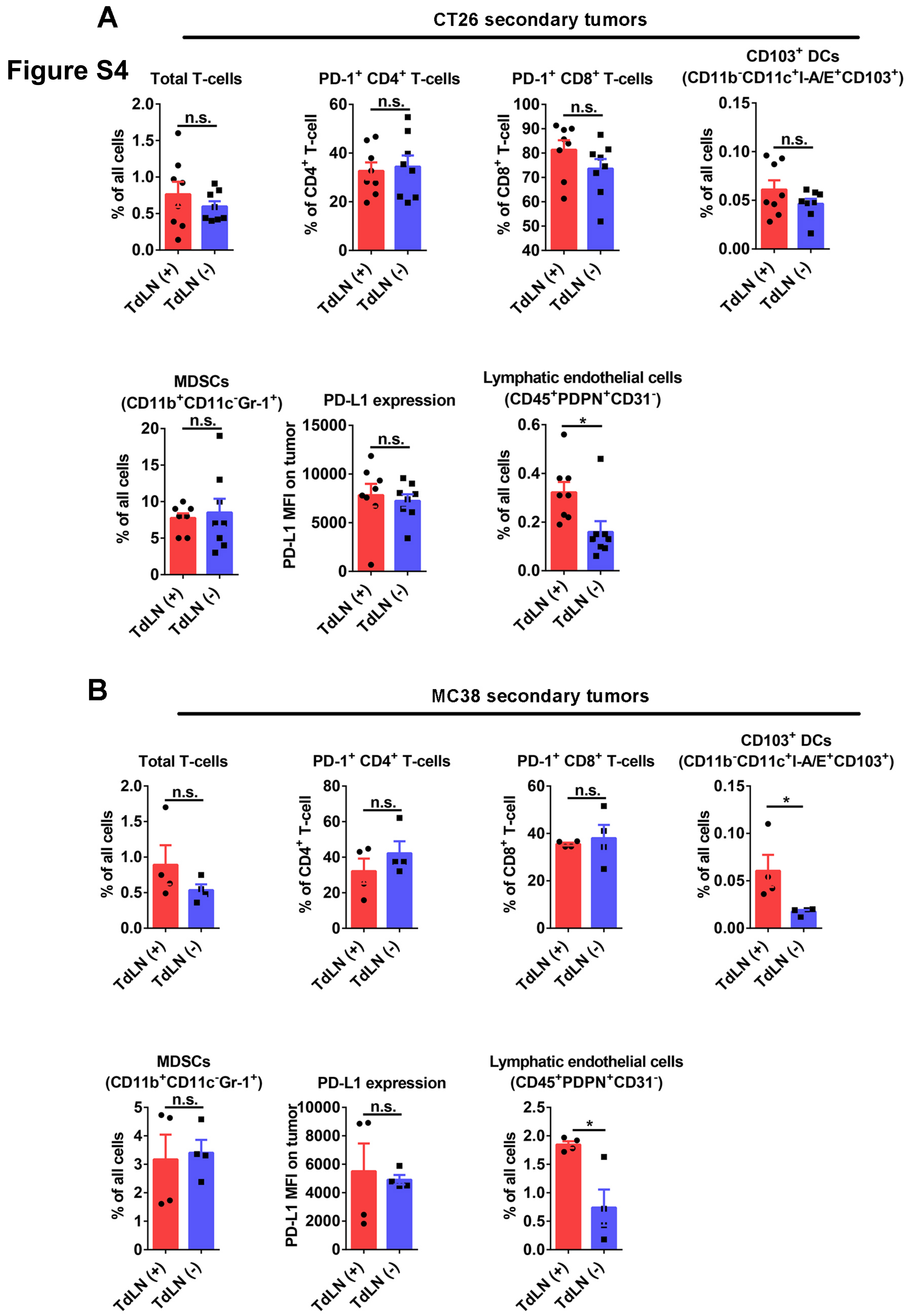
Figure S4
A
CT26 secondary tumors
Total T-cells
PD-1+ CD4+ T-cells
PD-1+ CD8+ T-cells
CD103+ DCs
(CD11b-CD11c+I-A/E+CD103+)
MDSCs
(CD11b+CD11c-Gr-1+)
PD-L1 expression
Lymphatic endothelial cells
(CD45+PDPN+CD31-)
% of all cells
% of CD4+ T-cell
% of CD8+ T-cell
PD-L1 MFI on tumor
n.s.
*
TdLN (+)
TdLN (-)
B
MC38 secondary tumors

**Figure S4. Immune features in secondary tumors with or without TdLNs. Related to Figure 2.**

**A)** The frequency of lymphatic endothelia cells was higher in CT26 secondary tumors (mimicking recurrent tumors) with TdLNs than secondary tumors without TdLNs (n=8 in each group, t-test). The total tumor-infiltrating T-cell frequency, PD-1 high expression T cells frequency, $CD103^+$ dendritic cells (DCs) frequency, myeloid-derived suppressive cells (MDSCs) frequency, and PD-L1 expression were similar in two groups (n=8 in each group, t-test was performed, data displayed as means ± SEMs, n.s.: no significance, *$p$<0.05).

**B)** The experiments were repeated in the MC38 tumor model. The frequency of lymphatic endothelial cells and $CD103^+$ DCs were higher in secondary tumors with TdLNs than in secondary tumors without TdLNs (n=4 in each group, t-test was performed, data displayed as means ± SEMs, n.s.: no significance, *$p$<0.05).

### Figure S5

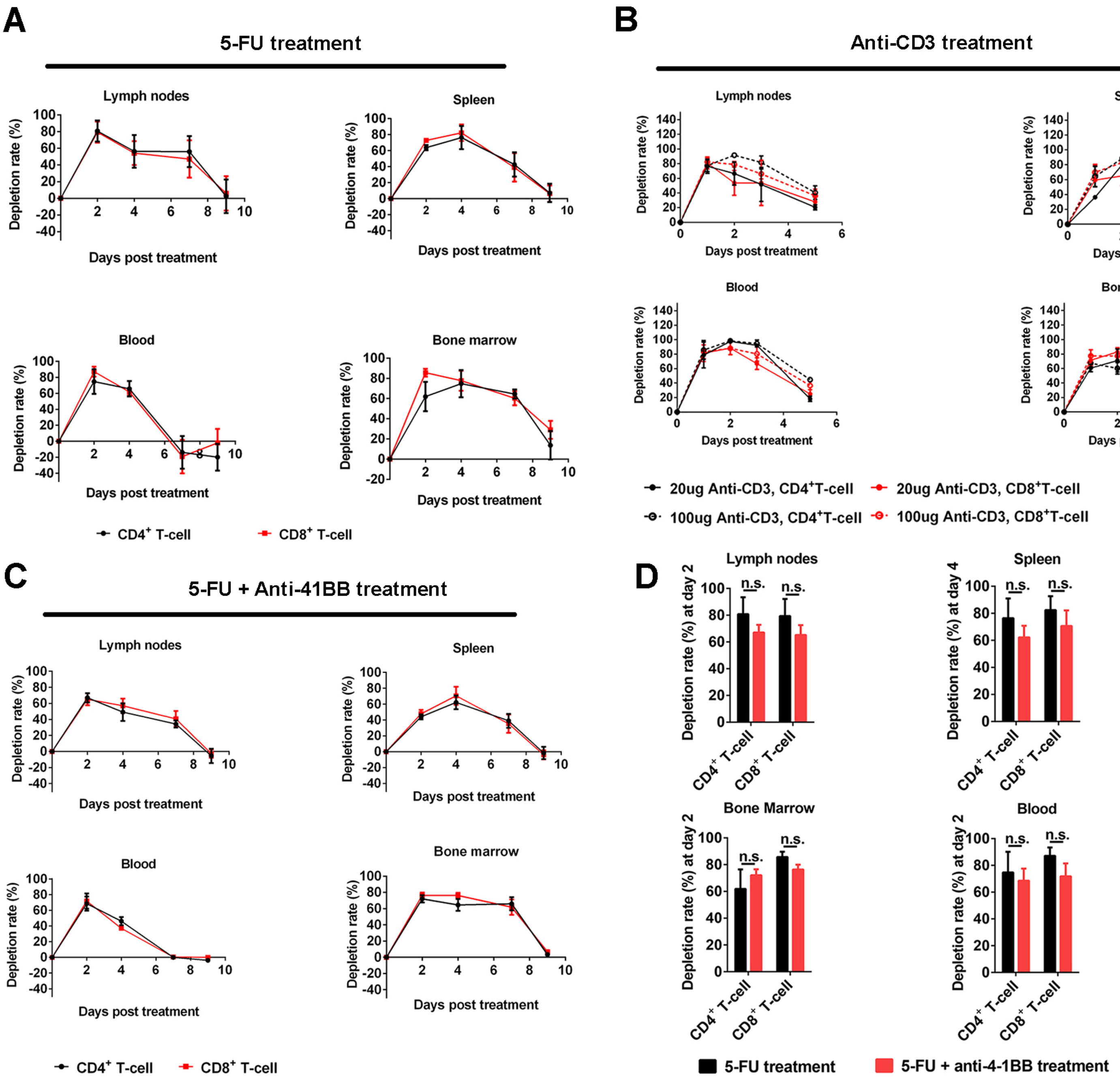
A
5-FU treatment
Lymph nodes
Spleen
Blood
Bone marrow
Depletion rate (%)
Days post treatment
CD4+ T-cell
CD8+ T-cell
B
Anti-CD3 treatment
Lymph nodes
Spleen
Blood
Bone marrow
Depletion rate (%)
Days post treatment
20ug Anti-CD3, CD4+T-cell
20ug Anti-CD3, CD8+T-cell
100ug Anti-CD3, CD4+T-cell
100ug Anti-CD3, CD8+T-cell
C
5-FU + Anti-41BB treatment
Lymph nodes
Spleen
Blood
Bone marrow
Depletion rate (%)
Days post treatment
CD4+ T-cell
CD8+ T-cell
D
Lymph nodes
Spleen
Bone Marrow
Blood
Depletion rate (%) at day 2
Depletion rate (%) at day 4
n.s.
CD4+ T-cell
CD8+ T-cell
5-FU treatment
5-FU + anti-4-1BB treatment

**Figure S5. T-cell depleting effects of 5-FU and anti-CD3 treatment. Related to Figure 2 and 4-6.**
**A)** 5-FU treatment on naïve mice depleted T cells in lymphatic organs, blood circulation, and bone marrow. The T-cell population was recovered around 9 days after 5-FU treatment (n=3 in each group, data were displayed as means ± SEMs).
**B)** Single-dose of anti-CD3 treatment on naïve mice depleted T cells for around 3 days (n=3 in each group, data were displayed as means ± SEMs).
**C-D)** A combination of anti-4-1BB with 5-FU didn't rescue the T-cell depletion induced by 5-FU treatment (n=3 in each group, t-test was performed, data were displayed as means ± SEMs, n.s.: no significance).

### Figure S6

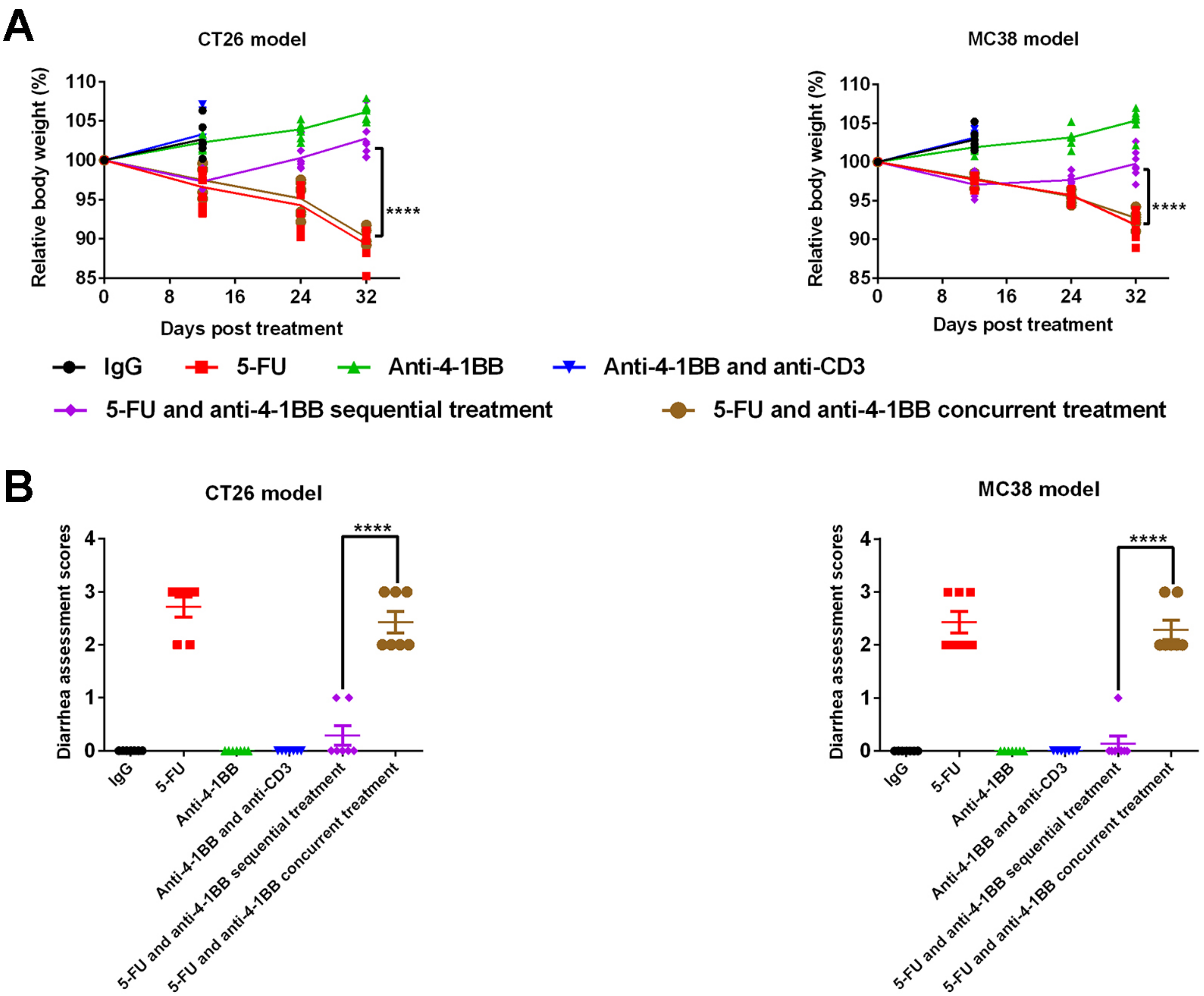

A
CT26 model
MC38 model
Relative body weight (%)
Days post treatment
****
IgG
5-FU
Anti-4-1BB
Anti-4-1BB and anti-CD3
5-FU and anti-4-1BB sequential treatment
5-FU and anti-4-1BB concurrent treatment
B
CT26 model
MC38 model
Diarrhea assessment scores
****
IgG
5-FU
Anti-4-1BB
Anti-4-1BB and anti-CD3
5-FU and anti-4-1BB sequential treatment
5-FU and anti-4-1BB concurrent treatment

**Figure S6. Side effects of different treatments. Related to Figure 4.**

**A-B)** The mouse body weight was measured on day 12, 24, and 32 during treatment (related to figure 4). On day 32, the mice treated with the 5-FU and anti-4-1BB sequential treatment have higher body weight than the mice treated with the 5-FU and anti-4-1BB concurrent treatment (n=7 in each group, t-test was performed between indicated groups at the last time point, individual value was shown, ****$p$<0.0001).

**C-D)** Diarrhea assessment was performed at the endpoint of mice follow-up to evaluate the side effects on mouse intestine. The 5-FU and anti-4-1BB concurrent but not sequential treatment caused severe diarrhea (n=7 in each group, t-test was performed between indicated groups, data were displayed as means ± SEMs, ****$p$<0.0001).

**Figure S7**

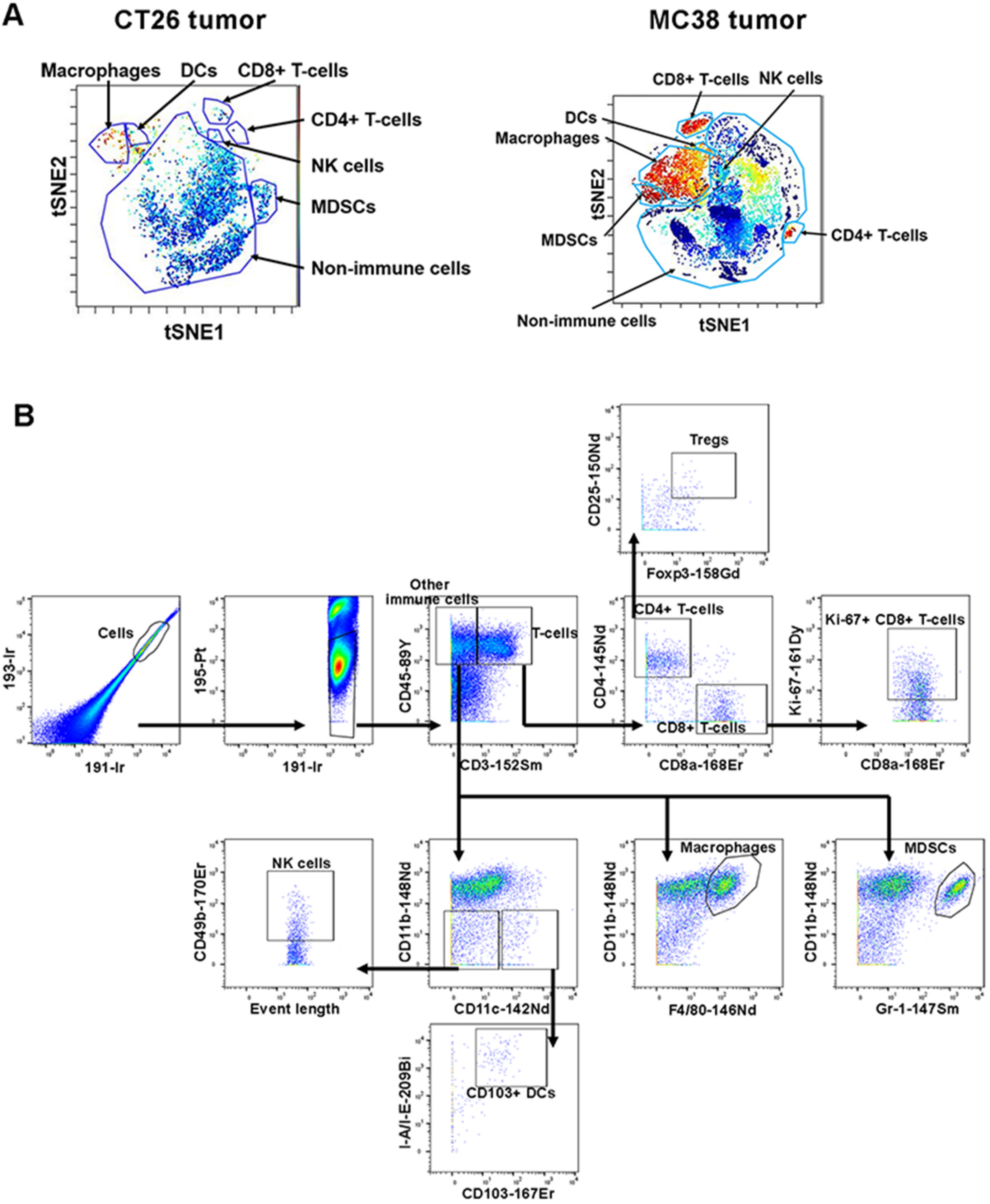

A
CT26 tumor
Macrophages
DCs
CD8+ T-cells
CD4+ T-cells
NK cells
MDSCs
Non-immune cells
tSNE2
tSNE1
MC38 tumor
CD8+ T-cells
NK cells
DCs
Macrophages
MDSCs
CD4+ T-cells
Non-immune cells
tSNE2
tSNE1
B
Tregs
CD25-150Nd
Foxp3-158Gd
Cells
193-Ir
191-Ir
195-Pt
191-Ir
Other immune cells
T-cells
CD45-89Y
CD3-152Sm
CD4+ T-cells
CD8+ T-cells
CD4-145Nd
CD8a-168Er
Ki-67+ CD8+ T-cells
Ki-67-161Dy
CD8a-168Er
NK cells
CD49b-170Er
Event length
CD11b-148Nd
CD11c-142Nd
Macrophages
CD11b-148Nd
F4/80-146Nd
MDSCs
CD11b-148Nd
Gr-1-147Sm
CD103+ DCs
I-A/I-E-209Bi
CD103-167Er

**Figure S7. Gating of the tumor-infiltrating immune cells. Related to Figure 6.**
**A)** The major tumor-infiltrating immune cell populations were showed in the tSNE plots.
**B)** Manual gating of the major tumor-infiltrating immune cells. The alive cell population was first identified and the immune cells (CD45+) were then gated. The gating of T-cell populations ($CD45^+CD3^+CD8^+$ for $CD8^+$ T cells, $CD45^+CD3^+CD4^+CD25^+Foxp3^+$ for $T_{regs}$), NK cells ($CD45^+CD3^-CD11b^-CD11c^-CD49b^+$), macrophages ($CD45^+CD3^-CD11b^+F4/80^+$), myeloid-derived suppressive cells (MDSCs, $CD45^+CD3^-CD11b^+Gr\text{-}1^+$), and $CD103^+$ DCs ($CD45^+CD3^-CD11b^-CD11c^+I\text{-}A/I\text{-}E^+CD103^+$) were showed here. The same markers were used throughout the study to identify the immune cells.

### Figure S8

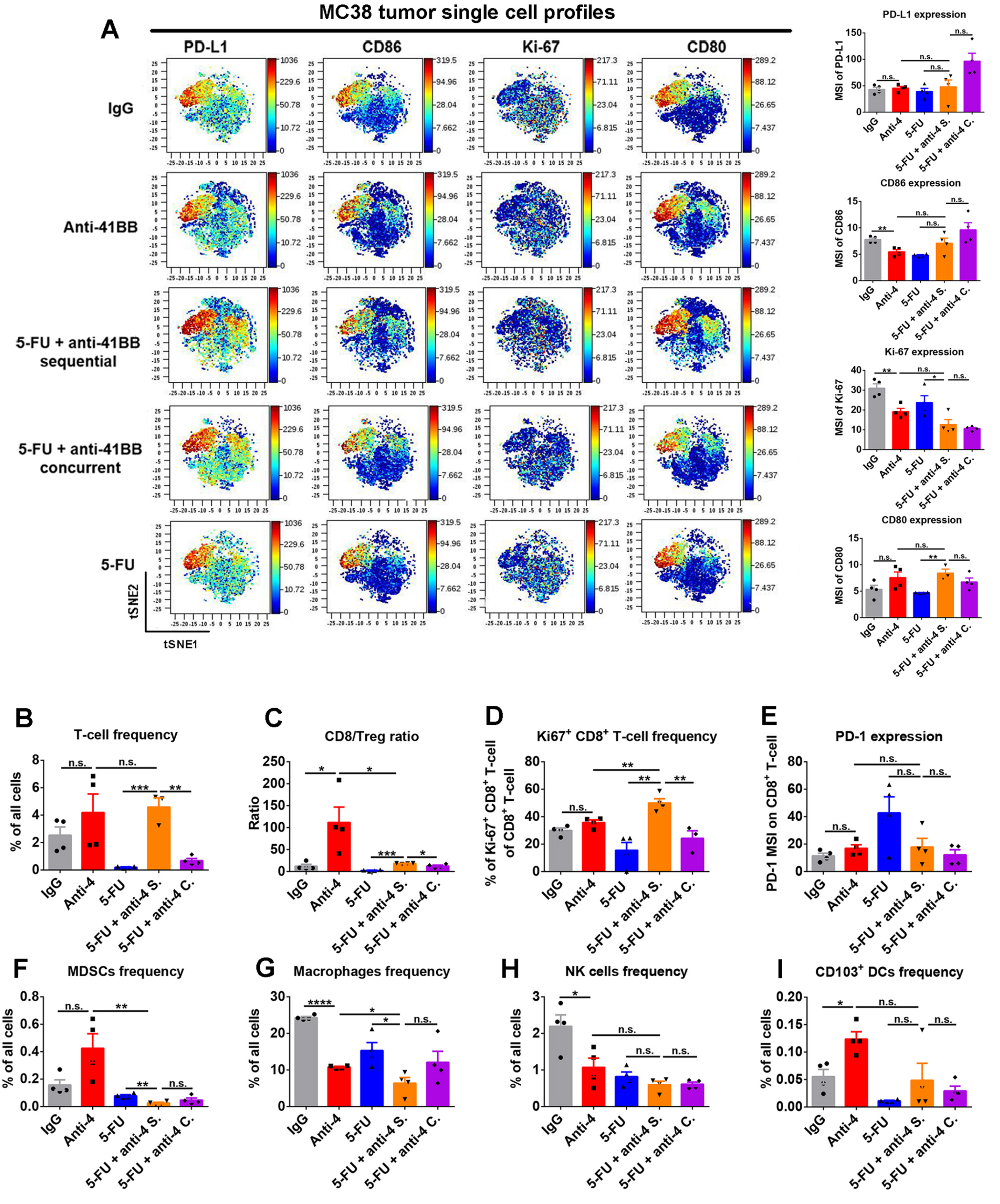

A
MC38 tumor single cell profiles
PD-L1
CD86
Ki-67
CD80
IgG
Anti-41BB
5-FU + anti-41BB sequential
5-FU + anti-41BB concurrent
5-FU
tSNE2
tSNE1
PD-L1 expression
CD86 expression
Ki-67 expression
CD80 expression
B
T-cell frequency
C
CD8/Treg ratio
D
Ki67+ CD8+ T-cell frequency
E
PD-1 expression
F
MDSCs frequency
G
Macrophages frequency
H
NK cells frequency
I
CD103+ DCs frequency
IgG
Anti-4
5-FU
5-FU + anti-4 S.
5-FU + anti-4 C.

**Figure S8. Tumor immunological response to 5-FU and anti-4-1BB treatments in MC38 tumors. Related to Figure 6.**

**A)** ViSNE plot showed single cell level expression of PD-L1, Ki-67, CD80, and CD86 in the MC38 tumor tissue. The 5-FU and anti-4-1BB sequential treatment significantly upregulated CD80 while decreased Ki-67 expression in tumor tissues (n=4 in each group, t-test was performed between indicated groups, data were displayed as means ± SEMs, MSI: mean signal intensity, n.s.: no significance, *$p$<0.05, **$p$<0.01).

**B-I)** The tumor-infiltrating T-cell frequency, CD8/Treg ratio, and Ki-67$^+$ CD8$^+$ T-cell frequency were higher in the sequential treatment than the concurrent treatment group. The myeloid-derived suppressive cells (MDSCs) were depleted in 5-FU treated groups. The 5-FU and anti-4-1BB sequential treatment were compared with the anti-4-1BB monotherapy, 5-FU monotherapy, and 5-FU and anti-4-1BB concurrent treatment (n=4 in each group, t-test was performed between indicated groups, data were displayed as means ± SEMs, MSI: mean signal intensity, n.s.: no significance, *$p$<0.05, **$p$<0.01, *** $p$<0.001, ****$p$<0.0001).

### Figure S9

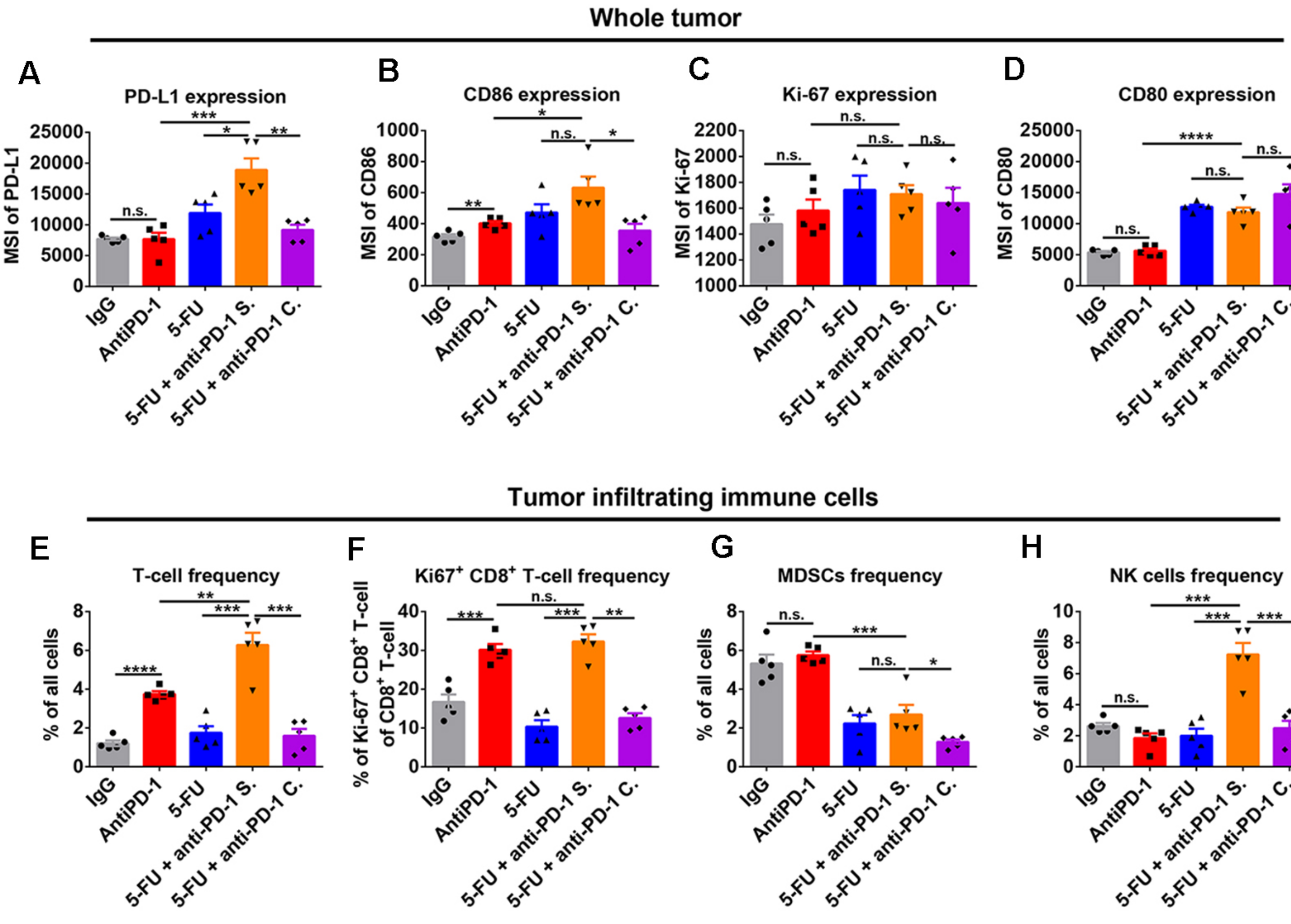

Whole tumor
A
PD-L1 expression
MSI of PD-L1
B
CD86 expression
MSI of CD86
C
Ki-67 expression
MSI of Ki-67
D
CD80 expression
MSI of CD80
IgG
AntiPD-1
5-FU
5-FU + anti-PD-1 S.
5-FU + anti-PD-1 C.
Tumor infiltrating immune cells
E
T-cell frequency
% of all cells
F
Ki67+ CD8+ T-cell frequency
% of Ki-67+ CD8+ T-cell of CD8+ T-cell
G
MDSCs frequency
% of all cells
H
NK cells frequency
% of all cells

**Figure S9. Tumor immunological response to 5-FU and anti-PD-1 treatments in CT26 tumors. Related to Figure 6.**

**A-D)** The 5-FU and anti-PD-1 sequential treatment significantly increased PD-L1, CD80, and CD86 expression in CT26 tumors (n=5 in each group, t-test was performed between indicated groups, data were displayed as means ± SEMs, MSI: mean signal intensity, n.s.: no significance, *$p$<0.05, **$p$<0.01, *** $p$<0.001, ****$p$<0.0001).

**E-H)** The 5-FU and anti-PD-1 sequential treatment significantly increased tumor-infiltrating T-cell frequency, Ki-67$^{+}$CD8$^{+}$ T-cell frequency, and NK cell frequency in tumor tissues, compared with the concurrent treatment. The myeloid-derived suppressive cells (MDSCs) were depleted in 5-FU treated groups (n=5 in each group, t-test was performed between indicated groups, data were displayed as means ± SEMs, MSI: mean signal intensity, n.s.: no significance, *$p$<0.05, **$p$<0.01, *** $p$<0.001, ****$p$<0.0001).

**Table S1. gp70 mRNA expression in different tissues. Related to Figure 1-3.**

| **Model** | **Tumor tissue** | **Tumor marginal skin & connective tissues post-surgery** | **Normal skin tissue** |
|---|---|---|---|
| **CT26 (Babl/c)** | Positive | Negative | Negative |
| **MC38 (C57BL/6)** | Positive | Negative | Negative |

**Transparent Methods**

*Cell cultures*

Murine CRC cell lines CT26 (purchased from American Type Culture Collection (ATCC)) and MC38 (gift from Dr. Nicholas Haining) were used for the study and were authenticated by STR profiling. CT26 cells were maintained in complete RPMI-1640 medium (GIBCO BRL), supplemented with 10% heat-inactivated FBS (Thermo Fisher Scientific), 100 IU/mL penicillin, and 100 μg/mL streptomycin (Invitrogen Life Technologies). MC38 cells were cultured in the complete DMEM medium (GIBCO BRL) with the same supplements as the RPMI 1640 medium. All cells were routinely authenticated and tested for mycoplasma.

*Mice*

Wild type BALB/c mice (6-8 weeks old, Jackson Laboratory) and C57BL/6 mice (6-8 weeks old, Charles River Laboratories) were used for animal studies. All mice were kept in a specific pathogen-free facility with fully autoclaved cages to minimize non-tumor specific immune activation. Animal studies were approved by the institutional animal care and use committee (IACUC). All mice are female. We don't expect the any influence of gender on our study aims.

*Subcutaneous tumor induction*

For the subcutaneous syngeneic model, the cells were harvested at low passages, washed, and resuspended in Matrigel matrix (Corning Inc.) before injection. Mice were shaved right before injection. CT26 ($2\times10^5$ cells/injection) or MC38 ($5\times10^5$ cells/injection) cells were inoculated subcutaneously into the right hind-flank of 6 to 8-week-old female BALB/c or C57BL/6 mice. The same amount of tumor cells were used for the rechallenge experiment in Figure 1. Tumor length and width were measured every three to seven days, and the volume was calculated according to the formula (length $\times$ width$^2$)/2. Mice were divided into different experimental groups at random when tumors reached a specific size.

*Identification of major tumor-draining lymph nodes*

To identify the major tumor-draining lymph nodes (TdLNs), we injected 50μl 1% Evans blue (Sigma-Aldrich) or 50μl 1% Alexa Fluor® 488 dye (Thermo Fisher Scientific) into the subcutaneous tumor (~400-500mm$^3$) at the right hind flank. The left and right inguinal LNs, axillary LNs, brachial LNs, popliteal LNs, and mesentery LNs were taken at 10 min, 30 min, and 60 min post Evan blue injection. For the fluorescence-labeled group, we collected LNs at 0.5h, 3h, 24h, and 48h post-injection. The intact LNs were visually examined for Evans blue staining. LNs, spleen, and tumor tissues were ground and meshed for single cell suspension, which was measured by flow cytometry for Alexa Fluor® 488 dye signal. To evaluate the physical change of LNs and spleen during tumor development, we weighted LNs and spleen from naïve mice and mice with different sizes of the tumor (100-200mm$^3$, 500-700mm$^3$, or 1200-1500mm$^3$).

*Subcutaneous tumor and TdLNs resection*

Primary tumors were resected when they reached the indicated volume as shown in the experimental schematics in each figure. Tumor-bearing mice were anesthetized with Ketamine (100 mg/kg) and Xylazine (10 mg/kg) by intraperitoneal injection. To minimize animal pain, we administrated Buprenorphine (slow-releasing, 2 mg/kg) subcutaneously 2 hours before anesthesia. Mice were prepared by removing hair from the skin region over the tumor. We prepared the skin by wiping with iodine prep pads and then alcohol prep pads. Resections were performed by elliptical incisions, 5mm left to the subcutaneous tumors. With iris scissors, we separated the capsule of subcutaneous tumors from the surrounding connective tissue to isolate and resect intact tumors. Once tumors were removed from the adjacent fascia, the incisions were sutured with 5/0 vicryl ties (polyglactin 910, Ethicon). For the TdLNs resection, the TdLNs were located based on the superficial anatomic landmark points. The mice were prepared as mentioned above. A 5-10mm incision was made and TdLNs were removed. Then the skin was sutured with 5/0 vicryl ties. For tumor rechallenge, 1 day after surgery, we inoculated the secondary tumor (CT26: $5\times10^5$ cells/injection, MC38: $1\times10^6$ cells/injection) to the surgical site to mimic tumor recurrence.

*RT-qPCR*

We used the murine leukemia virus envelope gp70 as a biomarker of tumor burden. Biopsies were collected from normal mouse skin, tumor tissue, and surgical margin after tumor resection. The mirVana microRNA (miRNA) Isolation Kit (Thermo Fisher Scientific) was used to extract total RNA from these biopsies. 500 ng of total RNA was used for establishing the cDNA library with the QuantiTect Reverse Transcription Kit (Qiagen). We used the LightCycler 480 Instrument (Roche Life Science) to measure 18S ribosomal RNA (rRNA) and gp70 expression.

Primers used: 18S rRNA forward primer: GTTGGTTTTCGGAACTGAGG, 18S rRNA reverse primer: AGTCGGCATCGTTTATGGTC, gp70 forward primer: AAAGTGACACATGCCCACAA, gp70 reverse primer: CCCCAAGAGGCACAATAGAA(Scrimieri et al., 2013).

*Flow cytometry*

Flow cytometry was used to measure tumor tissue immune infiltration, tumor antigen-specific T cells, and immune cell functions. Harvested tumor tissues were chopped into small pieces (around 3mm x 3mm) and then digested in a solution of collagenase IV (1 mg/ml) and deoxyribonuclease (DNase, 50 units/ml) at 37°C for 1 hr with shaking. The digested tissue was then meshed and filtered through a 70 μm cell strainer. The cell suspension was centrifuged and resuspended in red blood cell lysis buffer for 15 minutes at room temperature for eliminating red blood cells. Another centrifugation was performed to get the cell pellet for staining. For the lymphatic organs, we directly meshed the tissue and filtered through a 40 μm cell strainer to get single cell suspension, followed by red blood cells elimination.

Following the tissue sample preparation, cells were stained with the fixable cell viability dye and then cell surface marker antibodies for a 15 min incubation at 4°C. Next, cells were fixed and permeabilized for intracellular staining for a 30 min incubation at RT. The cells were finally stained with intracellular markers (30 min at RT) and analyzed on a BD FACS-CANTO instrument (BD Biosciences). To analyze the tumor antigen-specific T cells, we performed H-2Ld MuLV gp70-SPSYVYHQF APC conjugated tetramer (MBL International) staining by following the manufacturer's instruction, before antibody staining. The influenza hemagglutinin-IYSTVASSL APC conjugated tetramer (MBL International), which should only stain a very minimal population of T cells in mice without influenza hemagglutinin stimulation, was used as a negative control for ruling out false positive in the tetramer staining and setting up the gate for gp70 tetramer. Lymphatic tissues from naïve mice were also used as negative controls. According to the manufacture's instruction and our preliminary experiment optimization, we used anti-CD8 (clone KT15) antibody (MBL International) to further reduce false-positive rate of the tetramer staining. All antibodies for flow cytometry were purchased from Biolegend and summarized in supplementary materials. Data were analyzed using FlowJo software (Tree Star, Inc.).

*Mass cytometry*

Details on antibodies and reagents used are listed in supplementary table 2. We purchased the prelabeled antibodies from Fluidigm Corporation and unlabeled antibodies (MaxPar® Ready purified) from Biolegend. Conjugation of the purified antibodies with metal tags was performed by using the MaxPar X8 antibody labeling kit (Fluidigm Corporation) according to the manufacturer's instructions. The metal tagged antibodies were then validated and titrated in positive control and negative control samples.

Tumor samples were collected and digested using standard flow cytometry procedure. A total of 3 million single cells were used for each mass cytometry staining. In brief, the single cell pellets were first incubated with Cell-ID Cisplatin with a final concentration of 5 μM for 5 min at RT to identify dead cells. Cells were then washed and blocked by Fc-receptor blocking solution. Cell membrane staining was then performed with metal-conjugated antibodies for 30 min at RT. After staining, cells were fixed and permeabilized. The intracellular staining antibodies were then added and incubated for 45 min at RT. Finally, cells were labeled with 1 ml 1,000× diluted 125 μM Cell-ID intercalator-Ir to stain all cells in MaxPar Fix and Perm Buffer overnight at 4 °C. EQ Four Element Calibration Beads with the reference EQ passport P13H2302 were added to each staining tube right before data acquisition by a CyTOF 2 mass cytometer. The mass cytometry data were then normalized and exported for gating on alive single cells, which were then imported to the Cytobank software. A t-SNE analysis was performed with default parameters (perplexity, 30; iterations, 1,000) on all cell types in tumor samples.

*Mouse IFN-γ enzyme-linked immunosorbent assays*

Mouse naïve lymph nodes and TdLNs were collected, weighed, and ground in 100 μl RIPA lysis and extraction buffer. After the tissues were lysed, the total protein was used for enzyme-linked immunosorbent assay (ELISA, Affymetrix) to detect mouse *IFNγ*, by following the manufacturer's protocol.

*Histology*

Mouse naïve lymph nodes, TdLNs, and non-tumor draining lymph nodes (NdLNs) were collected and fixed in 10% formalin for 24 hr. Tissues were embedded in paraffin and cut for hematoxylin and eosin (H&E) staining. The whole tissue sections were scanned and analyzed for potential metastatic tumor cells.

*T-cell depletion*

We tested the effects of 5-FU treatment and 5-FU and anti-4-1BB combination treatment on T-cell depletion *in vivo*. Intraperitoneal administration of anti-CD3 treatment (clone: 17A2, BioXcell, 5 mg/kg every 3 days) was given to induce T-cell depleted mice. One dose of 5-FU (150 mg/kg) or 5-FU (150 mg/kg) and anti-4-1BB (5 mg/kg) combination treatment was given intraperitoneally in naïve mice. Mice were sampled on days 2, 4, 7, and 9 after treatment for quantifying T cells in lymph nodes, spleen, bone marrow, and blood circulation.

*Mouse treatments*

Mice were treated with IgG (5 mg/kg as an anti-4-1BB control, 10 mg/kg as an anti-PD-1 control), 5-FU (150 mg/kg), anti-4-1BB agonist (5 mg/kg, clone: 3H3), or anti-PD-1 (10 mg/kg, clone: RMP1-14) for treatment purpose. For the 5-FU monotherapy, one dose of 5-FU was given every 12 days to minimize the severe side effects. For anti-4-1BB and IgG monotherapy, mice were treated every 3 days. For the 5-FU and anti-4-1BB sequential treatment, anti-4-1BB treatment started 9 days after one dose 5-FU treatment and continued as 3 days per injection after that. For the 5-FU and anti-4-1BB concurrent treatment, we added the anti-4-1BB cycle to the 5-FU cycle. The anti-PD-1 was used as the same as the anti-4-1BB cycle. All treatments were given intraperitoneally and continued until the endpoint of study design. The treatment starting points and endpoints varied in different experiments for different purposes and were shown in the individual figure or figure legend.

*5-FU toxicity evaluation*

We recorded animal body weight and diarrhea scores after treatments. Mice were weighed on day 12, 24, and 32 after treatment. The diarrhea score was assessed at the endpoint of each treatment by using a 4-point scoring system: 0=normal stool; 1=slight diarrhea (soft formed stool without perianal staining of the coat); 2=moderate diarrhea (unformed stool with moderate perianal staining of the coat); and 3=severe diarrhea (watery stool with severe perianal staining of the coat)(Song et al., 2013).

*Statistical analysis*

All statistical analyses and graphing were performed using GraphPad Prism software (Version 6). Data were displayed as means ± SEMs. For comparison of two groups quantitative data, paired or unpaired Student's t-test was performed. When applicable, one-way analysis of variance (ANOVA) was utilized for multiple groups' comparison, followed by post hoc (Tukey's) multiple comparisons test. Kaplan-Meier curves were plotted to visualize mouse survival, and log-rank tests were used to compare survival outcomes between subgroups. A two-tail P value of less than 0.05 was considered statistically significant.

**Table of key materials**

| **Reagent for immune assays** | **Clone** | **Vendor** | **Identifier** |
|---|---|---|---|
| Anti-mouse CD3-FITC | 17A2 | BioLegend | 100204 |
| Anti-mouse CD28-PE | 37.51 | BioLegend | 102106 |
| Anti-mouse PD-1-PerCP/Cy5.5 | 29F.1A12 | BioLegend | 135208 |
| Anti-mouse CD62L-PE/Cy7 | MEL-14 | BioLegend | 104418 |
| Anti-mouse CD8a-APC/Cy7 | 53-6.7 | BioLegend | 100714 |
| Anti-mouse LAG-3-BV421 | C9B7W | BioLegend | 125221 |
| Anti-mouse/human CD44-PE | IM7 | BioLegend | 103008 |
| Anti-mouse CD19-Pacific Blue | 6D5 | BioLegend | 115523 |
| Anti-mouse CD4-BV510 | GK1.5 | BioLegend | 100449 |
| Anti-mouse CD86-PE | GL-1 | BioLegend | 105007 |
| Anti-mouse F4/80-PE/Cy5 | BM8 | BioLegend | 123111 |
| Anti-mouse CD80-PE/Cy7 | 16-10A1 | BioLegend | 104734 |
| Anti-mouse/human CD11b-APC | M1/70 | BioLegend | 101212 |
| Anti-mouse I-A/I-E-APC/Cy7 | M5/114.15.2 | BioLegend | 107628 |
| Anti-mouse CD11c-BV510 | N418 | BioLegend | 117338 |
| Anti-mouse CD45-Pacific Blue | 30-F11 | BioLegend | 103126 |
| Anti-mouse CD45-FITC | 30-F11 | BioLegend | 103108 |
| Anti-mouse CD3-PerCP/Cyanine5.5 | 17A2 | BioLegend | 100218 |
| Anti-mouse CD3ε-PE/Cy7 | 145-2C11 | BioLegend | 100320 |

| | | | |
|---|---|---|---|
| Anti-mouse CD103-Pacific Blue | 2E7 | BioLegend | 121418 |
| Anti-mouse Gr-1-PE/Cy7 | RB6-8C5 | BioLegend | 108416 |
| Anti-mouse CD45-BV510 | 30-F11 | BioLegend | 103138 |
| Anti-mouse Podoplanin-APC/Cy7 | 8.1.1 | BioLegend | 127418 |
| Anti-mouse CD31-Pacific Blue | 390 | BioLegend | 102422 |
| Anti-mouse CD45-89Y | 30-F11 | Fluidigm | 3089005B |
| Anti-mouse Ly-6G-41Pr | 1A8 | Fluidigm | 3141008B |
| Anti-mouse CD11c-142Nd | N418 | Fluidigm | 3142003B |
| Anti-mouse CD4-145Nd | RM4-5 | Fluidigm | 3145002B |
| Anti-mouse F4/80-146Nd | BM8 | Fluidigm | 3146008B |
| Anti-mouse Gr-1-147Sm | RB6-8C5 | BioLegend/Fluidigm | 108449/201147B |
| Anti-mouse CD11b-148Nd | M1/70 | Fluidigm | 3148003B |
| Anti-mouse CD19-149Sm | 6D5 | Fluidigm | 3149002B |
| Anti-mouse CD25-150Nd | 3C7 | Fluidigm | 3150002B |
| Anti-mouse CD28-151Eu | 37.51 | Fluidigm | 3151005B |
| Anti-mouse CD3e-152Sm | 145-2C11 | Fluidigm | 3152004B |
| Anti-mouse CD274-153Eu | 10F.9G2 | Fluidigm | 3153016B |
| Anti-mouse CD152-154Sm | UC10-4B9 | Fluidigm | 3154008B |
| Anti-mouse CD279-155Gd | RMP1-30 | BioLegend/Fluidigm | 109113/201155A |
| Anti-mouse CD335-156Gd | 29A1.4 | BioLegend/Fluidigm | 137625/201156B |
| Anti-mouse Foxp3-158Gd | FJK-16s | Fluidigm | 3158003A |
| Anti-mouse RORgt-B2D-159Tb | B2D | Fluidigm | 3159019B |
| Anti-mouse CD62L-160Gd | MEL-14 | Fluidigm | 3160008B |
| Anti-mouse Ki-67-161Dy | B56 | Fluidigm | 3161007B |
| Anti-mouse Ly-6C-162Dy | HK1.4 | Fluidigm | 3162014B |
| Anti-mouse CD197-164Dy | 4B12 | Fluidigm | 3164013A |
| Anti-mouse IFNg-165Ho | XMG1.2 | Fluidigm | 3165003B |
| Anti-mouse IL-4-166Er | 11B11 | Fluidigm | 3166003B |
| Anti-mouse CD103-167Er | 2E7 | BioLegend/Fluidigm | 121402/ 201167B |
| Anti-mouse CD8a-168Er | 53-6.7 | Fluidigm | 3168003B |
| Anti-mouse CD49b-170Er | HMa2 | Fluidigm | 3170008B |
| Anti-mouse CD80-171Yb | 16-10A1 | Fluidigm | 3171008B |
| Anti-mouse CD86-172Yb | GL1 | Fluidigm | 3172016B |
| Anti-mouse Granzyme B-173Yb | GB11 | Fluidigm | 3173006B |
| Anti-mouse CD127-174Yb | A7R34 | Fluidigm | 3174013B |
| Anti-mouse CD44-176Yb | IM7 | BioLegend/Fluidigm | 103051/201176B |
| Anti-mouse I-A/I-E-209Bi | M5/114.15.2 | Fluidigm | 3209006B |
| Cell-ID™ Intercalator-Ir | NA | Fluidigm | 201192B |
| Cell-ID™ Cisplatin | NA | Fluidigm | 201064 |
| H-2Ld MuLV gp70 Tetramer-APC | NA | MBL International | TB-M521-2 |
| IFN gamma Mouse ELISA Kit | NA | Thermo Fisher Scientific | BMS606 |
| Zombie Violet™ Fixable Viability Kit | NA | BioLegend | 423113 |
| Zombie Aqua™ Fixable Viability Kit | NA | BioLegend | 423101 |
| Zombie Green™ Fixable Viability Kit | NA | BioLegend | 423111 |
| MuLV gp70 Tetramer-APC | NA | MBL International | TB-M521-2 |
| Influenza HA Tetramer-APC | NA | MBL International | TS-M520-2 |
| Anti-mouse CD8-FITC | KT15 | MBL International | D271-4 |
| **Drugs or antibodies for treatment** | **Clone** | **Vendor** | **Identifier** |
| Anti-mouse 4-1BB (CD137) | 3H3 | BioXcell | BE0239 |
| Anti-mouse PD-1 | RMP1-14 | BioXcell | BP0146 |
| Anti-mouse CD3 | 17A2 | BioXcell | BE0002 |
| 5-Fluorouracil | NA | Intas Pharmaceuticals | DB00544 |

| | | | |
|---|---|---|---|
| **Mouse** | **Age** | **Vendor** | **Identifier** |
| BALB/cJ | 6-8 weeks | The Jackson Laboratory | 000651 |
| C57BL/6 | 6-8 weeks | Charles River | 027 |
